# Individual beliefs about temporal continuity explain variation of perceptual biases

**DOI:** 10.1101/2021.07.13.452167

**Authors:** Stefan Glasauer, Zhuanghua Shi

## Abstract

Perception of magnitudes such as duration or distance is often found to be systematically biased. The biases, which result from incorporating prior knowledge in the perceptual process, can vary considerably between individuals. The variations are commonly attributed to differences in sensory precision and reliance on priors. However, another factor not considered so far is the implicit belief about how successive sensory stimuli are generated: independently from each other or with certain temporal continuity. The main types of explanatory models proposed so far – static or iterative – mirror this distinction but cannot adequately explain individual biases. Here we propose a new unifying model that explains individual variation as combination of sensory precision and beliefs about temporal continuity and predicts the experimentally found changes in biases when altering temporal continuity. Thus, according to the model, individual differences in perception depend on beliefs about how stimuli are generated in the world.

## Introduction

Magnitude estimates are pervasive in our daily activities, such as predicting forthcoming events, estimating the travel distance, and precisely controlling our movements. However, we also make perceptual errors. Some of these errors have been found to be systematic biases, which have been investigated throughout the history of psychophysics. Two perceptual biases, the central tendency effect (Vierordt 1868; Hollingworth 1910) and the sequential dependence (Holland & Lockhead 1968; Cross 1973), are still hotly debated till today (Shi et al. 2013; Fischer & Whitney 2014; Petzschner et al. 2015). The central tendency effect refers to a systematic overestimation of small magnitudes and underestimation of large magnitudes, whereas the sequential dependence (or ‘serial dependence’, Fischer & Whitney 2014) designates that the current perceptual estimate depends not just on the current stimulus, but also on stimuli given in the past. While both biases have long been accepted as inevitable properties of magnitude perception, several quantitative theoretical accounts only emerged in the last decade that linked them to perception being a form of Bayesian inference (Jazayeri & Shadlen 2010; Petzschner & Glasauer 2011; Cicchini et al. 2012; Roach et al. 2017). The concept of Bayesian inference also provides an operational explanation for interindividual differences seen in perception: Bayesian inference considers a perceptual estimate to result from a near-optimal combination of sensory inputs and prior knowledge. The extent to which sensory input and prior knowledge are weighted depends on the accuracy of sensory measurement and the certainty of prior knowledge. When driving through fog, prior knowledge of the road ahead is more important than on a sunny day. Thus, under the Bayesian framework, individual sensory accuracy and precision about the prior knowledge (formalized as width of the prior distribution) is commonly thought to explain inter-individual differences (Petzschner & Glasauer 2011; Powell et al. 2016). For example, individual differences in color perception may reflect different properties of the eye and retina (sensor) or contextual influences such as the experienced environment (prior) (Mollon et al. 2017).

However, Bayesian perception offers an alternative account of individual differences: the prior knowledge that is used to improve the sensory input depends on our implicit assumptions about how sensory stimuli are caused, or generated, in the external world. These assumptions, or beliefs, are essential for predictions, which serve as prior knowledge: for example, if asked what tomorrow’s temperature will be, we might answer something like ‘a little warmer/colder than today’. That is, we assume that daily temperature changes by a random amount each day, but that it will be similar on successive days. In contrast, in a standard psychophysical experiment the stimuli presented to our participants are often drawn randomly from a fixed distribution, just like numbers in a lottery, and they are thus independent from trial to trial. Thus, a good guess (in the sense of small error) of predicting the value presented at the next trial would be the mean of the stimulus distribution. In other words, when it comes to daily temperature, a good estimator would use today’s temperature as prior knowledge for tomorrow’s value. In contrast, when the stimulus value presented in a psychophysical experiment is concerned, a useful prior knowledge would be the distribution of possible values.

Thus, when a sensory input is combined with prior knowledge using these two cases of generative assumptions, the final estimate might differ substantially (Fig. 1)^1^. Hence, two individuals that have such different beliefs about how sensory events are, or are not, linked over time, may exhibit different perceptual biases. Bayesian theory suggests this possibility, but to our knowledge this has not yet been investigated.

**Figure 1:**
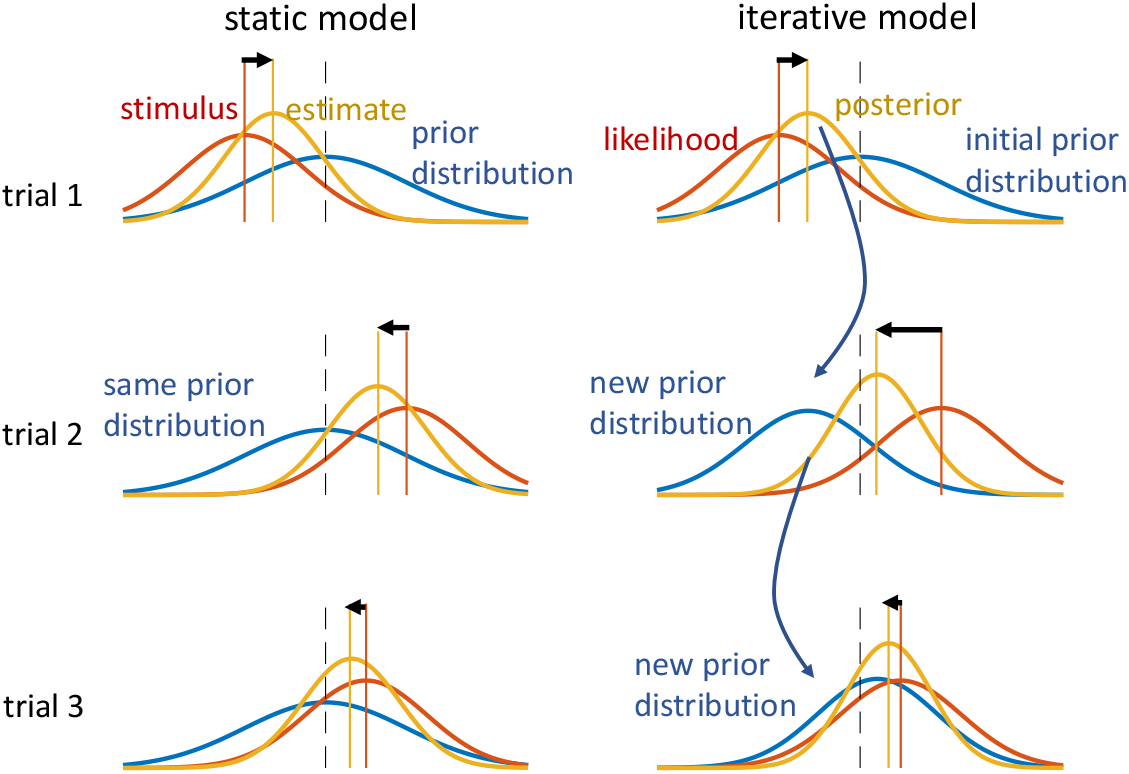
Two estimation models (static and iterative) for magnitude perception. The static model uses static time-independent prior knowledge (blue prior distribution), while the iterative model dynamically updates the prior knowledge in each trial. In trial 1 (upper row), in both cases the prior knowledge (blue) and the stimulus (orange line), to which a likelihood distribution (orange) is attached, are the same for both models. Thus, the estimate, shown as posterior distribution (yellow) and computed by multiplying prior and likelihood, is also equal for both cases, already showing the characteristic overestimation of small amplitudes. The black dashed line serves as visual reference and is aligned with the mean of the initial prior. In trial 2 (middle row), the stimulus and likelihood are again the same for both models, but the prior differs: the static model takes the same prior as in trial 1, because the underlying assumption is that all stimuli come from that distribution. In contrast, the iterative model uses the estimate from the previous trial to predict the next stimulus distribution, which is used as new prior knowledge, because it assumes that the new stimulus is similar to the old except for some random change. The two resulting posterior distributions (yellow), and thus the perceptual estimates, differ considerably with the one for the static model showing much smaller underestimation than that of the iterative estimate, even though the stimuli and sensory accuracy is the same in both cases. The third row shows trial 3, in which the estimates of both models come much closer again.

Both cases, however, are mirrored by explanatory perceptual estimation models found in the literature. For example, Jazayeri and Shadlen (2010) proposed a model to account for the central tendency in duration reproduction in which prior knowledge was formalized as fixed stimulus distribution (as depicted in Fig. 1 as static model). By contrast, Petzschner & Glasauer (2011) in a distance reproduction study proposed that the central tendency is a consequence of an iterative Bayesian process, in which prior knowledge is iteratively based on the perceptual estimate of the previous trial (like in the iterative model in Fig. 1), rather than assuming a fixed prior. In the following years, various other similar explanations have been proposed, some of them using iterative updating (Dyjas et al. 2012; Thurley 2016), others assuming static priors (Cicchini et al. 2012; Roach et al. 2017). The common idea of these models is that the central tendency bias is a by-product of optimizing the perceptual estimates using prior knowledge when sensory input is uncertain.

While both types of models can successfully explain the central tendency bias, they make very different predictions about sequential dependence, the second type of bias: only the iterative updating models predict sequential dependence for randomly distributed stimuli. In a model with static prior, the perceptual estimates are independent from trial to trial.

In the present paper, we show that experimental results from duration estimation and distance estimation show sequential dependence and thus favour iterative models of perceptual estimation. However, we demonstrate that neither the static nor the iterative mathematical models found in the current literature can fully explain the interindividual differences. We thus propose a Bayesian model that not only explains these differences, but also allows to predict individual behaviour in a novel experimental situation. Our results thus clearly support the idea that individual variations in perceptual biases are indeed a consequence of differing beliefs about the temporal continuity of stimulus generation.

## Results

We examined central tendency and sequential dependence in three experimental data sets of magnitude estimation: one from duration reproduction (Glasauer & Shi 2019, 2021; data published as Glasauer & Shi 2021b), and two from linear and angular distance reproduction (Petzschner & Glasauer 2011, data set published as Petzschner & Glasauer 2020).

The first dataset comes from a duration reproduction study (preliminary data were presented in Glasauer & Shi 2019, 2021; data are published as Glasauer & Shi 2021b). In the experiment, subjects (n=14, 7 female, average age 27.4) had to reproduce a visually presented stimulus duration by pressing and holding a key (see Fig. 2, inset). Each subject received a random sequence of 400 stimulus durations between 400 and 1900 ms. We quantified both the sequential dependence and the central tendency effect using simple linear regression (see Materials and Methods for the detailed method). Fig. 2 shows an example of individual raw results plotted for evaluation of central tendency (Fig. 2A) and sequential dependence (Fig. 2B). The relation of sequential dependence and central tendency for all individual participants is depicted in Fig. 2C. Note that, as mentioned in the introduction, perceptual estimation with a fixed prior (see the static model in Fig. 1) would predict that sequential dependence is zero independently of central tendency.

**Figure 2:**
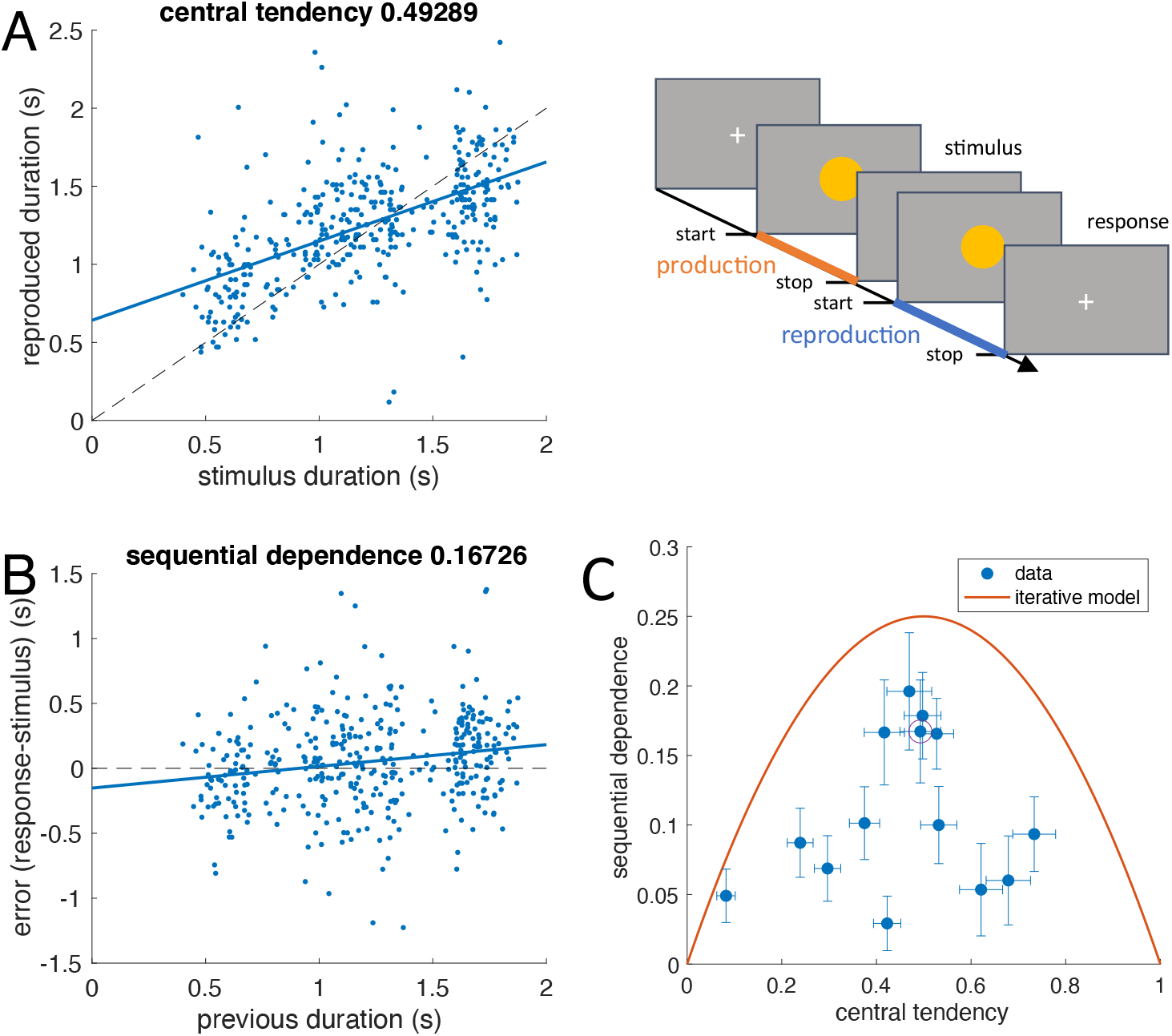
Central tendency and sequential dependence in a duration reproduction experiment. Inset: schematic procedure of a single trial. A visual cue is shown on a screen to indicate stimulus duration (production phase). Reproduction is performed by pressing a button (see also Glasauer & Shi 2019, 2020, and Materials and Methods for details). **A** & **B**: Raw data (blue dots) from one subject (400 trials). Stimuli were presented in random order. **A**: Reproduced duration plotted over stimulus duration; the central tendency is shown by the regression line (blue). **B**: Error (response-stimulus) in the current trial plotted over the stimulus duration of the previous trial. The sequential dependence is shown by the blue line. **C**: Sequential dependence plotted over central tendency for all subjects (n=14). Error bars denote standard error of the mean. The data point resulting for the single subject data presented in A and B is indicated by a purple circle. The orange parabola shows the predicted relation for the iterative model depicted in Fig. 1 (see SI Appendix A3: Serial dependence for an iterative Bayesian model).

However, individual responses show a large scatter for both central tendency and sequential dependence. The mean sequential dependence was 0.108 ± 0.056 (mean ± SD), which is significantly different from zero (*p* < .0001; *t*-test, n = 14) and thus ruled out the static model as valid explanation for the results. In fact, all data points show higher sequential dependence than predicted by the static model (zero). We also conducted a partial correlation analysis and calculated the correlation coefficient between current error and previous stimulus after controlling for the current stimulus. The average partial correlation coefficient was 0.197 ± 0.068 and significantly different from zero (*p* < .0001; *t*-test, n = 14). For comparison, the corresponding partial correlation coefficient between error and current stimulus, after controlling for the previous stimulus, was −0.623 ± 0.156 (*p* < .0001; *t*-test, n = 14), again revealing the central tendency.

As mentioned before, the sequential dependence larger than zero rules out the static prior as an explanation for the central tendency. However, in the iterative model (Fig. 1), sequential dependence is expected to occur, because the last posterior distribution is used as new prior in the next trial, thus introducing a temporal dependence across trials. When using this updating procedure (Glasauer 2019; see Materials & Methods for the model definition), the model predicts a fixed parabola relation between central tendency and serial dependence, shown in Fig. 2C (for a derivation see SI Appendix A3). However, individual serial dependence not only shows values higher than zero (the prediction of the static model), but also lower than the prediction of the iterative model. The average difference between the observed sequential dependence and that predicted by the iterative model was significant, (mean ± SD: −0.112 ± 0.053, *p* < .0001; *t*-test, *n* = 14), showing that also the iterative model cannot adequately predict the sequential dependence. It should be noted that the steady state of the iterative updating is equivalent to the “internal reference model” (Dyjas et al. 2012, Bausenhart et al. 2014) proposed for duration estimation.

We also analysed a publicly available data set on distance reproduction (Petzschner & Glasauer 2020) published previously (Petzschner & Glasauer 2011), using the same method. The data come from two separate experiments on visual path integration, one on linear distance reproduction and one on reproduction of angular distance (see Materials and Methods). While Petzschner and Glasauer (2011) showed that their iterative model could well capture the central tendency, they did not analyse sequential dependence. Fig. 3 shows the equivalent analysis as above for the two path-integration experiments (Petzschner & Glasauer 2011). For the linear distance reproduction, the average sequential dependence is 0.100 ± 0.045 (mean ± SD). For the angular distance reproduction, it is 0.119 ± 0.057 (mean ± SD). As for duration reproduction, the data for the two distance reproduction experiments confirms that neither the static nor the iterative model can capture the sequential dependence sufficiently well (all *p*s < 0.0001).

**Figure 3:**
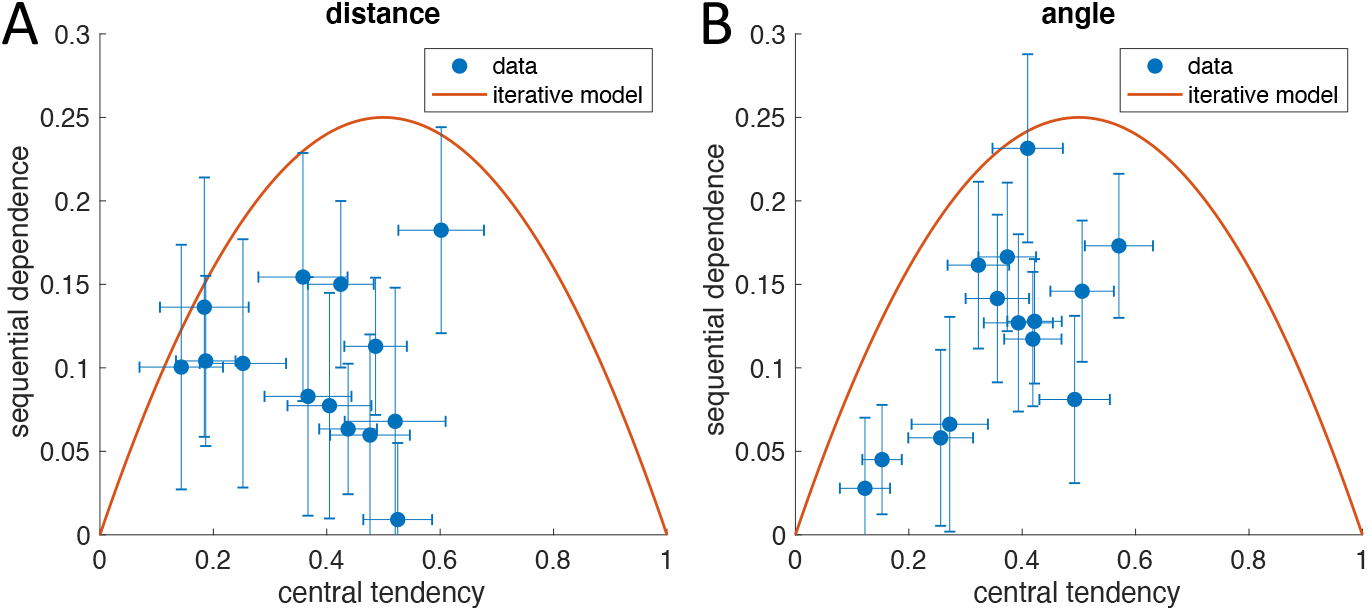
Sequential dependence for distance (**A**) and angular (**B**) reproduction plotted over central tendency for all subjects. Distance and angular reproduction data come from Petzschner & Glasauer 2011, data published as Petzschner & Glasauer 2020, each point corresponds to 540 trials. Error bars denote standard error of the mean. The same analysis was applied as in Fig. 2C.

The present data also show that the individual variation in biases cannot be explained by individually different levels of sensory uncertainty: under the assumption of a static model, changing sensory uncertainty would not lead to different levels of sequential dependence; if the iterative model was underlying perceptual estimation, variation of sensory uncertainty would still confine the individual biases to the orange line in Figs. 2C, 3A, and 3B.

From Figs. 2 and 3 we can see that for all individual data points the predictions of the two models considered so far seem to be boundary conditions. Neither is any of the individual bias values located below zero for sequential dependence, nor is any of them above the orange parabola denoting the iterative model. The obvious conclusion is that individual participants seem to follow beliefs that lie between the two extremes expressed by the static and simple iterative models, which assume 1) the sampled stimuli are random and independent, or 2) the current stimulus is equal to the previous plus some random change. An intermediate belief about stimulus generation can be described as follows: assume that the stimulus on the current trial has been drawn from a stimulus distribution, but that the mean of that distribution is allowed to change randomly from trial to trial. Regarding our first example of daily temperature changes, this assumption is also reasonably applicable.

From this assumption we can now construct a new estimation model, which requires two variables to be estimated: in addition to estimating the current stimulus, the perceptual process also needs to estimate the mean of the current stimulus distribution. Therefore, the model requires two internal states. The new model is an extended iterative model, more flexible than the simple iterative model depicted in Fig. 1. Since it comprises the static and iterative models of Fig. 1 as boundary conditions, the model is capable of simulating the individual biases of all three experiments (Fig 2C, 3A, 3B), but it comes with an increase of the number of free parameters from two to three (see Materials & Methods for model equations). To find out whether the new model is indeed superior in fitting the experimental data, we fitted^2^ the extended (two-state) iterative model (3 free parameters per subject) to the individual data of the duration experiment to evaluate whether the model would capture not only the central tendency but also the sequential dependence better than the simpler alternatives. An example time course of raw data (stimulus and response) together with best-fit simulation is shown in Fig 4A (see also SI Appendix C: fits to mean responses in Fig. S4A). Note that the fit minimizes the least-squares distance between the individual responses and the model simulation, with the models receiving as input the trial-to-trial time course of the stimuli in the same order as presented to the individual participant.

**Figure 4:**
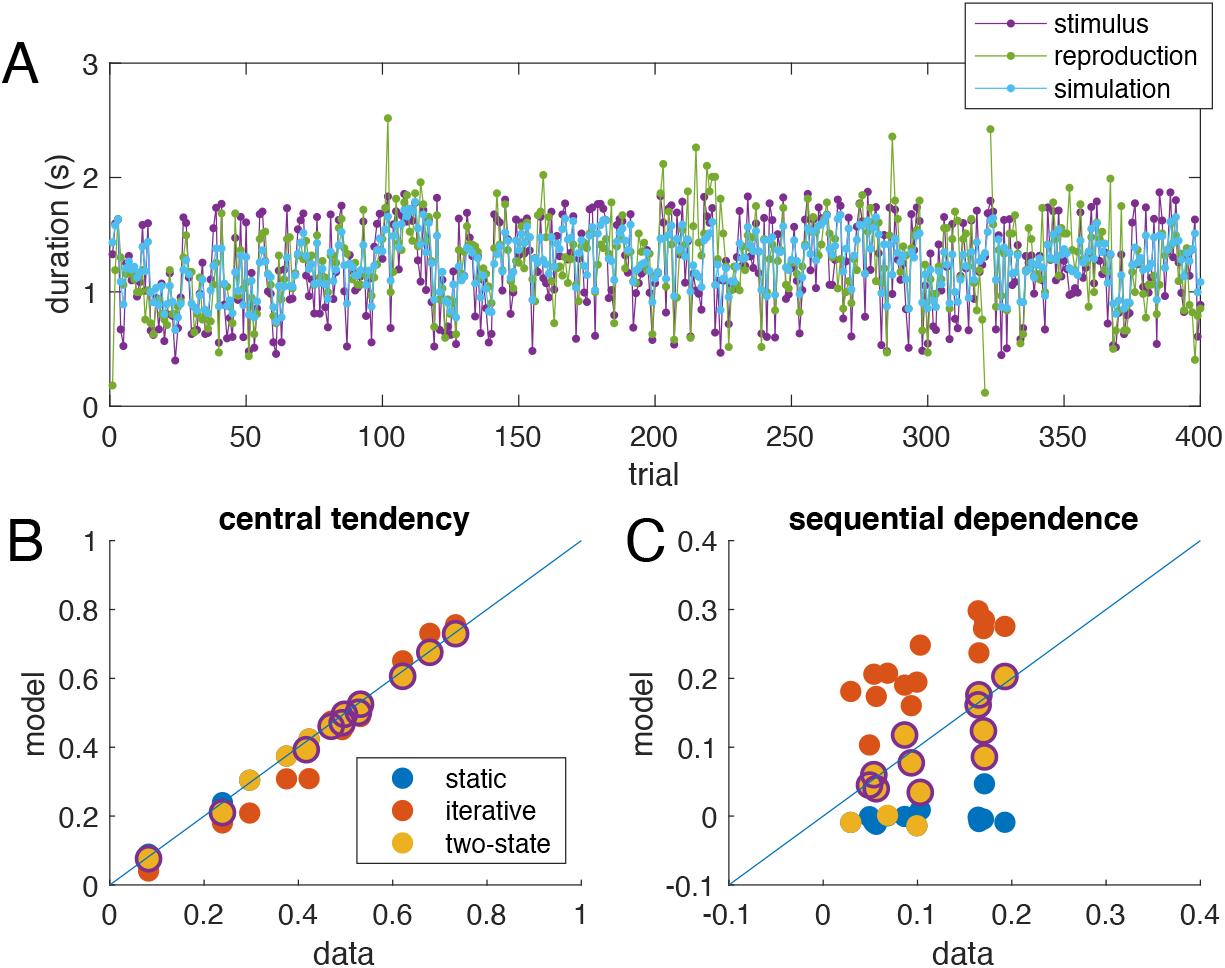
**A**: Time course of stimulus (magenta) and reproduction (green) from one participant (same data as in Fig. 2A,B) together with best fit of the two-state model (light blue). The model fitting minimized the least-squares distance between reproduction (green) and simulation (light blue). The input to the model was the trial-by-trial stimulus time course (magenta). The few outliers (e.g., around trial 105 or 320) were not removed for model fitting. **B, C**: Comparison of experimental data (x-axis) and model simulation (y-axis) for central tendency (**B**) and absolute sequential dependence (**C**) for all three models and all participants. Each dot represents one participant. The central tendency is captured correctly by all three models (values close to the diagonal), whereas the model values for sequential dependence are too small for the static model (blue dots, close to zero), and too large for the simple iterative model (red dots). Values for the two-state model lie closer to the diagonal in most cases. Cases for which the two-state model was the best model according to cross-validation are highlighted by circles: for 8 of 14 participants, the two-state model was superior, while in the remaining 6 cases the static model was sufficient to explain the data.

The static model and the simple iterative model are special cases of the new two-state model: both are nested within the two-state model. Therefore, one can determine whether the parameters that are set to zero for the simpler models differ significantly from zero in the full model. On average, both parameters (Parameter 1: the relative variability of the stimulus distribution, and Parameter 2: the relative variability of the additive change of the mean) of the full model were significantly different from zero (Parameter 1: 1.03 ± 0.28, mean ± SEM, *t*-test *p* < .01; Parameter 2: 0.14 ± 0.05, mean ± SEM, *t*-test p = 0.025; both *n* = 14). In individual participants, the relative variability of the stimulus distribution differed significantly from zero (assessed via confidence intervals of the parameters) for all subjects (range 0.20 to 4.12), while the variability of the additive change differed from zero only for 6 of 14 subjects (range 0 to 0.66). To determine which model was more appropriate for fitting the data, we used an out-of-sample cross validation procedure specifically suited for model selection in time series (Arlot & Celisse 2010). According to this cross-validation procedure (see Materials and Methods), the two-state model is the preferable model for 8 of 14 participants, while for the remaining 6 participants the static model is sufficient (Fig. 4B,C). A comparison of the values for central tendency and absolute sequential dependence derived from the data and from respective model simulations is presented in Fig 4B and 4C. In case of perfect fit, all points should lie on the diagonal. While all three models capture well the central tendency (Fig. 4B), the simulated serial dependence is too small for the static model but too large for the simple iterative model (Fig. 4C), while the two-state model matches the data reasonably well.

While the results so far confirm that the two-state model provides a quantitative explanation for central tendency and sequential dependence at lag one (i.e., dependence on the previous stimulus), due to its iterative nature the two-state model also predicts dependence of the current error on stimuli further in the past. That this is indeed the case also experimentally can be shown by cross correlation analysis: for duration reproduction, the cross correlation between stimulus and reproduction is, on average, significantly different from zero up to lag 3 (*t*-test; lag 2: *p* = 0.0007; lag 3: *p* = 0.039; *n* = 14; supplementary Fig. S5).

The averaged experimental results for duration reproduction together with the averaged model results for dependence of error on current and previous stimuli is shown in Fig. 5 (see also SI Fig. S4). Note that the model was fitted to each individual trial-by-trial reproduction time course separately by minimizing the trial-wise least-squares distance between experimental reproduction and model simulation. Thus, the good match shown in Fig. 5B and 5C, quantified by a high coefficient of determination R^2^, is caused by the model mimicking the experimental sequential dependence without explicitly including it in the fitting procedure. This is not a trivial consequence of the model fit, as shown by the fact that both static and simple iterative models can fit the central tendency equally well (i.e., the dependence shown in Fig. 5A), but fail to correctly exhibit the sequential dependence shown in Fig. 5B and 5C (see SI Appendix C1).

**Figure 5:**
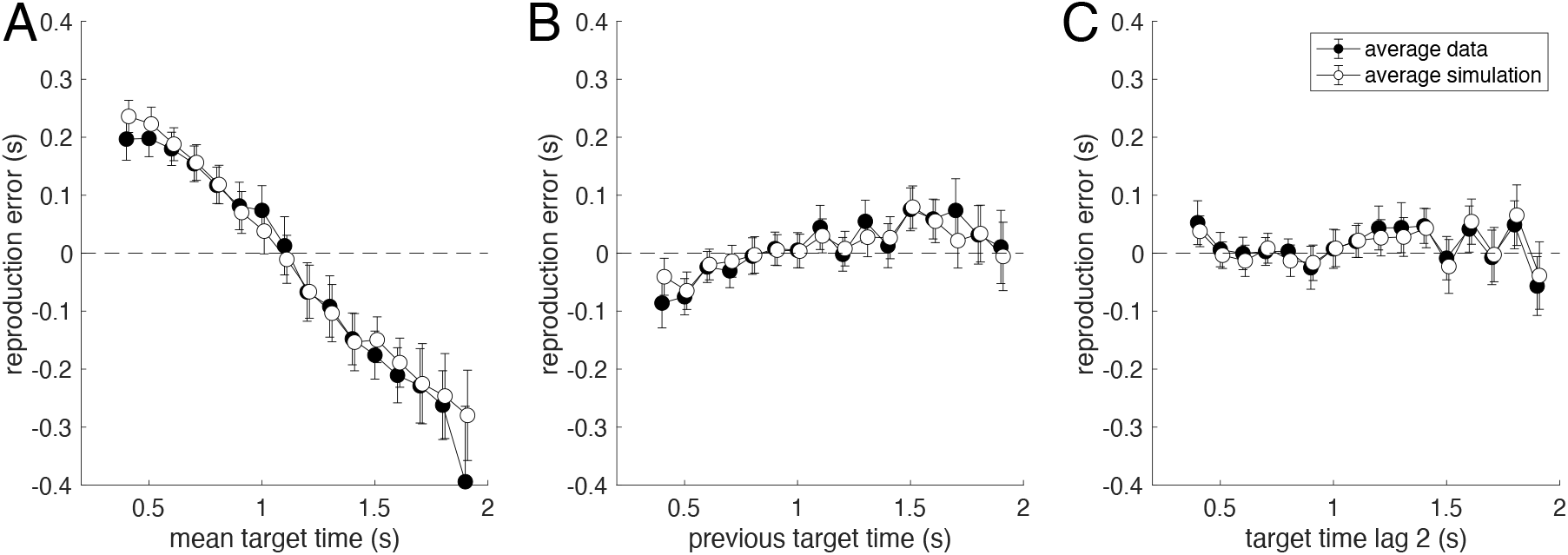
Average data for the duration reproduction experiment (black dots) together with average simulation result for the best fit iterative two-state model (open dots). Stimuli were presented in randomized order. **A**: Reproduction error as a function of the current target duration (R^2^=0.964). **B**: Sequential dependence shown as reproduction error plotted over previous target duration (i.e., R^2^=0.803). **C**: Reproduction error as a function of the target duration two trials in the past (R^2^=0.834). Data points are averages of 7 to 14 subjects (only 7 subjects received long durations above 1.7 s); error bars denote SEM. For the model simulation, the model (three free parameters) was fitted individually to the trial-by-trial time course of the responses of each subject (same fits as for Figure 4). From theses individual simulated response time courses, the average simulation was computed in the same way as for the real response time courses. Note that not only average mean data and simulated responses match, but that also the size of the error bars is captured quite well by the model.

As explained above, the models (static, iterative, two-state) make different assumptions about how stimuli are generated, e.g., the static model assumes independent and identically distributed (i.i.d.) random variables. Fig. 6 shows histograms, autocorrelation, and time course of exemplary stimulus sequences generated according to the assumptions of the three models. The stimulus sequences of the iterative models (iterative, two-state) have been generated so that their histograms are similar to the histogram of the static model’s sequence (quantified by minimizing the Kullback-Leibler divergence). While the histograms are reasonably similar (Fig. 6A, KL divergence < 0.01), the autocorrelation differs considerably (Fig. 6B). As expected, the sequence for the static model, which is a Gaussian noise sequence, shows no sequential dependence between current and previous values, while the two iterative models generate sequences with autocorrelation at higher lags. The sequence generated by the simple iterative model is a Wiener process or random walk, while the sequence of the two-state model is a superposition of a random walk and Gaussian noise. The corresponding exemplary time courses are shown in Fig. 6C: the blue time course (i.i.d. stimuli) would be optimal for the static model, the red random walk time course is optimal for the simple iterative model, and the yellow trace, being a compromise between randomness and slow drift would be optimal for the two-state model.

**Figure 6:**
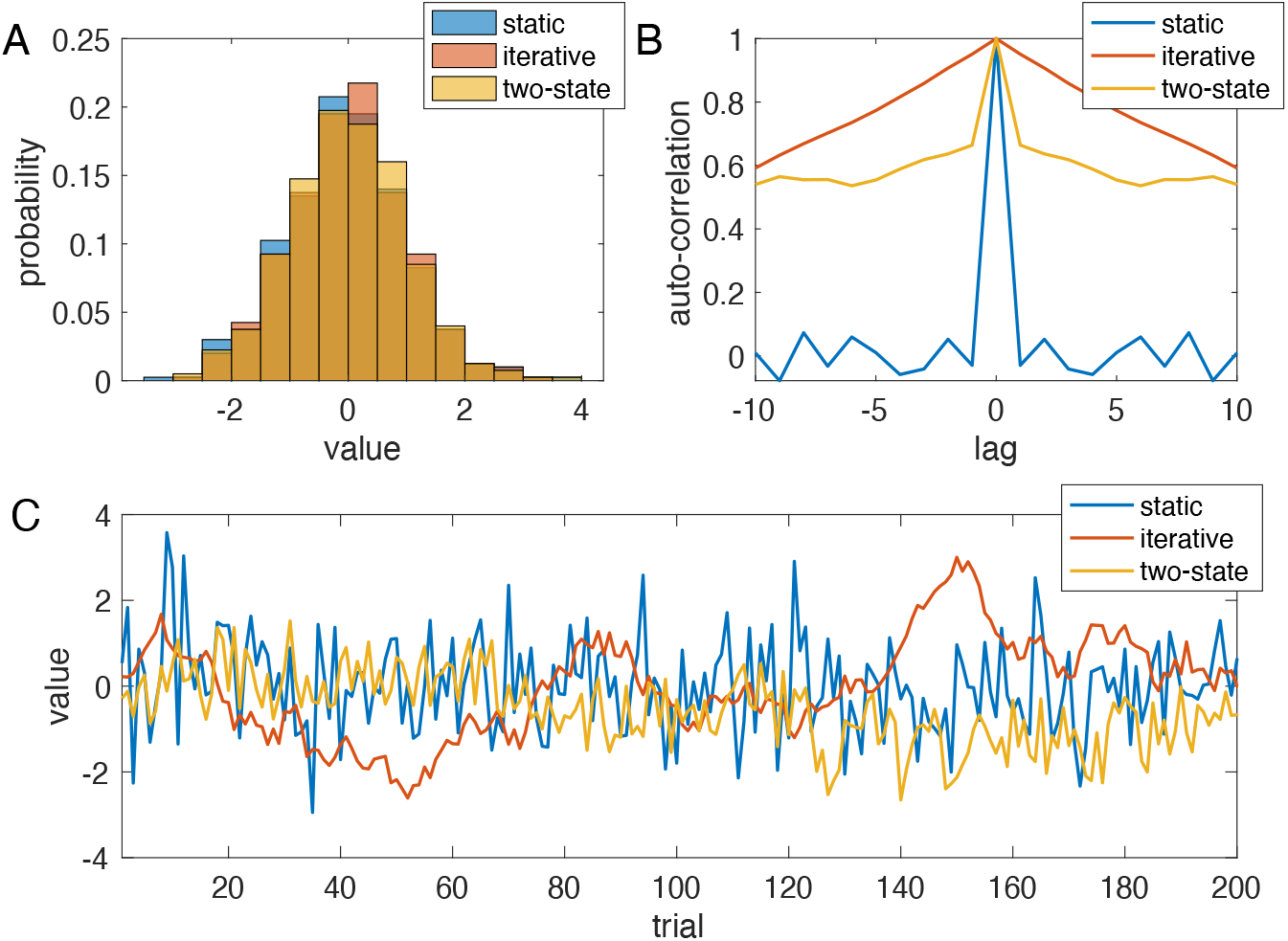
Sequences of stimuli generated by the three models. The sequences for the iterative and two-state model have been generated such that their histograms resemble that of the static sequence. **A**: histograms. **B**: autocorrelation. **C**: first 200 samples of all three sequences. The variances of the two-state model were chosen so that the autocorrelation at lag 1 equals approx. 0.66.

This implies that the result of the estimation process depends on how well the stimulus sequence is matched to the model assumptions about stimulus generation. Using an iterative model is suboptimal for an i.i.d. sequence, and, vice versa, using the static model is not the best solution for estimation a random walk sequence.

We tested this implication by analysing the second condition of the duration reproduction experiment, in which the same stimuli as analysed above, were presented in a random walk order to the same participants (see also Materials and Methods). Since we suspected that the remaining central tendency in the random walk condition and the change in sequential dependence could be explained by the new two-state model introduced above, we used the individually fitted model parameters obtained from the randomized condition to predict the individual time courses of the random walk condition. In this condition, subsequent stimuli are similar to each other (example in Fig. 7C), just as supposed by the generative model of the simple iterative Bayes (see Fig. 6C, red time course, for an example of such a random walk). As explained in our previous paper (Glasauer & Shi 2021), this condition tests the prediction of the simple iterative and the Petzschner & Glasauer (2011) explanatory models, which both predict that the central tendency vanishes in the random walk condition. An example of the effect of changed stimulus order on central tendency is shown in Fig. 7A (randomized condition, replotted from Fig. 2A). and Fig. 7B (random walk condition) for one participant. In this participant, the central tendency seen for randomized stimulus order (Fig. 7A) almost vanishes for the random walk stimulus order (Fig. 7B).

**Figure 7:**
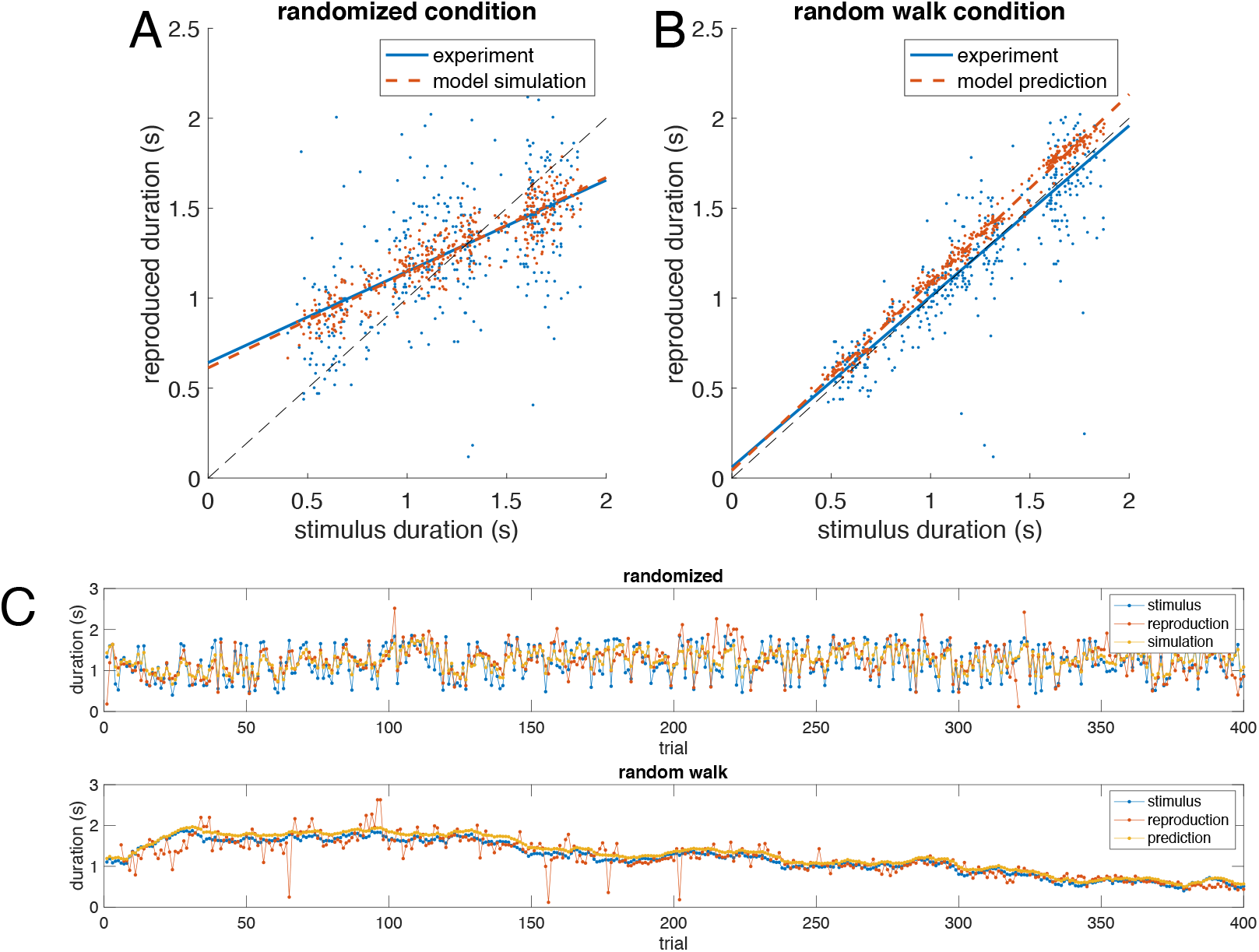
Example data from one participant and model results. **A**: Data (blue) and model simulation (best fit to two-state model, red) for the ‘randomized’ condition of the duration reproduction experiment. Same data as in Fig. 4A. Lines are linear least squares fits to the data showing that the model replicates the central tendency. **B**: Data (blue) and model *prediction* (red, model and parameters same as in A) for the ‘random walk’ condition, in which successive stimuli were similar. It can clearly be seen that in this subject the central tendency dramatically changed, which was predicted by the model. **C**: Data from A and B plotted over trial. The same stimuli were used in both conditions.

On average the data show that the central tendency indeed decreased substantially and was significantly smaller during random walk (*t*(13) = 7.32, *p* < .0001; see Fig. 8A). However, it did not completely vanish and was still larger than predicted by these previous models (Glasauer & Shi 2021). For some subjects, the central tendency was no longer different from zero (see example data in Fig. 7), while for others it clearly was still visible. Sequential dependence also changed and became on average negative with a significant difference between conditions (Fig. 8B; *t*(13) = 5.25, *p* < .001).

**Figure 8:**
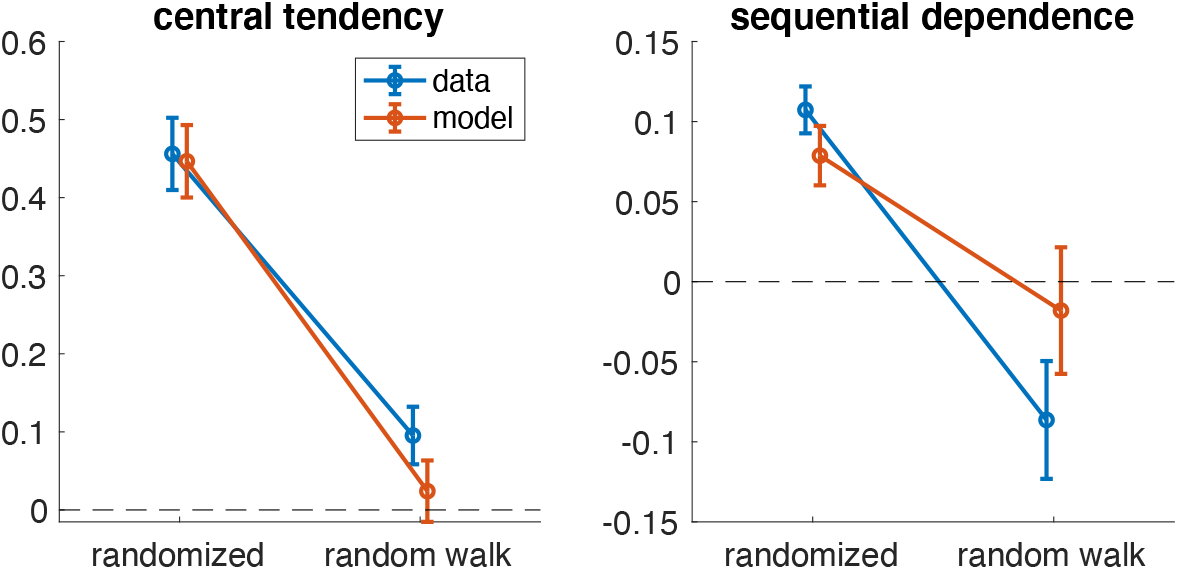
Summary of average results for both conditions of the duration reproduction experiment. Blue: data; orange: model. Error bars indicate standard error of the mean. **A:** Central tendency is significantly smaller during random walk (t-test, n = 14, p < 0.0001). **B**: Sequential dependence becomes on average negative with a significant difference between conditions (t-test, n=14, p = 0.00016). In A and B, the model values for the randomized condition are averages from the fitted simulation, while model values for the random walk condition are averaged *predictions* using the model parameters from the randomized condition.

Figure 9 shows the averaged experimental results together with the averaged model prediction. Both central tendency (Fig. 9A) and sequential dependence (Fig. 9B and 9C) are well-predicted by the model, showing that the central tendency remaining in the random walk condition is explained by the generative assumption of the two-state model (see also red model results in Fig. 8). Note that the similarity of the error dependence on current and previous stimuli in the random walk condition shown in Fig. 9 is expected, since stimuli in this condition are highly autocorrelated, i.e., the current stimulus is indeed similar to the stimuli preceding it (and thus the reproduction error is similar when plotted over current or previous stimuli).

**Figure 9:**
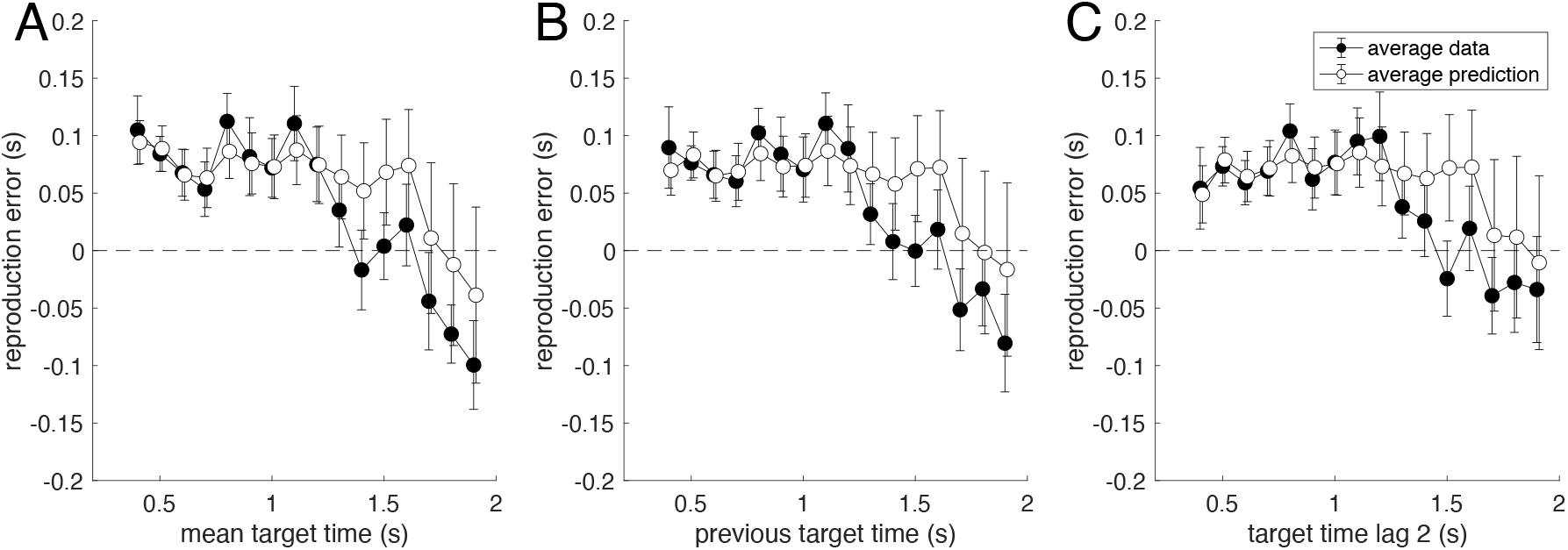
Average data for the ‘random walk’ condition of the duration reproduction experiment (black dots), in which successive stimuli were similar, presented together with averaged model predictions (not fits) of the iterative two-state models used in Fig. 5 (open dots). Stimuli are the same as in Fig. 5 except for the order of presentation. Model parameters were determined from fitting the ‘randomized’ condition and are the same as used in Fig. 5. **A**: Reproduction error as a function of the current target duration (R^2^=0.629). **B**: Reproduction error as a function of the previous target duration (R^2^=0.544). **C**: Reproduction error as a function of the target duration two trial in the past (R^2^=0.428). Data points are averages of 7 to 14 subjects (only 7 subjects received long durations above 1.7 s); error bars denote SEM. Average model predictions are calculated from trial-by-trial predicted individual time courses using the parameters corresponding to Fig. 5.

## Discussion

In this paper, we analysed the relation between the central tendency and sequential dependence for magnitude reproduction with two aims: to distinguish between static and iterative models proposed in the literature to explain the central tendency bias, and to reveal the origin of individual differences seen experimentally. We analysed three datasets, one from duration reproduction and two from path integration, to evaluate which model can better explain magnitude reproduction regarding both the sequential dependence, quantified as current error depending on previous stimulus magnitude, and the central tendency effects. Effects of immediate prior experience on current decisions have been reported for various cases in the psychological literature (e.g., Cross 1973, Fischer & Whitney 2014, Liberman et al. 2014), and, as we show here, they are also clearly visible in experiments on magnitude reproduction. The average sequential dependence found in duration and distance reproduction differed significantly from zero, which clearly demonstrates that the response error of the current trial depends on the stimulus from the last trial. This contradicts static models as explanation for perceptual biases because they imply no influence of the previous trial. Consequently, several previously published static trial-independent models can be ruled out (Jazayeri & Shadlen 2010, Roach et al. 2017, Lakshminarasimhan et al. 2018). Even though the static models fit the central tendency well in experiments with random stimulus presentation (see Fig. 4B), the explanation for this bias offered by the static models is only partially correct. The fundamental assumption of the static models, a fusion of the sensed stimulus with prior information about the stimulus range, is not completely wrong, except that the prior is not static, as shown by the significant sequential dependence. Rather, the prior is updated trial-by-trial so that information from the immediate previous trial is used for the current estimate. Due to the iterative nature of Bayesian estimation with the posterior as basis for the new prior, not just the previous stimulus (as proposed by Cicchini et al. 2018), but also stimuli further in the past can still exert an influence on the present response. This difference between static and iterative models has important consequences for understanding the processes that lead to the perceptual results: while the results of static and iterative models look similar with respect to the central tendency, the internally represented priors, the underlying assumptions about stimulus generation, and the predictions for the sequential dependence are completely different.

Like the static prior model, the simple iterative model proposed previously (e.g., Dyjas et al. 2012; Glasauer 2019) predicts the central tendency effect very well but falls short in accounting for the experimentally observed sequential dependence. The simple iterative model assumes that stimuli remain similar from trial to trial with a random fluctuation. This assumption corresponds to stimuli being generated by a random walk or discrete Wiener process. According to this assumption, the overall variance of the stimuli builds up over the trials. By contrast, the static model assumes that the stimulus distribution has a fixed variance, and a fixed mean. The generative assumption for the iterative model also implies a stimulus sequence that differs considerably from that of the static model: it resembles Brownian motion or a diffusion process in one dimension rather than a random sequence (see Fig. 6C for examples). Both the static and the simple iterative models provide predictions concerning the sequential dependence: the static model predicts zero sequential dependence, the iterative model predicts that, in case of random stimuli, sequential dependence depends on central tendency in a predictable way (see red curve in Figs. 2C and 3). The empirical data, however, showed that neither of these two models captures the experimental relation between central tendency and sequential dependence.

Therefore, we proposed the two-state model that combines the static and the simple iterative models and assumes that the stimulus at each trial comes from a distribution with fixed variance, but that the mean of that distribution changes from trial to trial. By merging the assumptions of static and simple iterative models about stimulus generation, both the central tendency effect and the absolute sequential dependence can be well explained. According to the two-state model, the considerable variations between participants are not only caused by different impact of noise on sensory measurement, but also because of different beliefs concerning the sequential structure of the stimuli. As an example, in Fig. 2C, of two participants with approximately the same central tendency of 0.42, one had a sequential dependence of 0.03, the other of 0.17. This difference reflects the observers’ own supposition about the sequential structure: the participant with a low sequential dependence assumed the world is volatile and trusted only the current stimulus together with a hypothesis about the limited range of stimuli for perceptual estimates. By contrast, the participant with a large sequential dependence agreed about the randomness of the world but further assumed that things change over time with some continuity. For perceptual decision-making task, it has recently been suggested that individual differences are due to different implicit assumptions about the complexity of a sequence (Glaze et al. 2018). In their study, participants had to infer from which of two possible Gaussian sources the current visual stimulus was drawn. The true source was randomly switched with a hazard rate that could change. The authors proposed that a bias-variance trade-off was the underlying reason for differences in choice variability. While this study is very different from ours, both have in common that the implicit beliefs of participants about the temporal volatility of stimulus generation are the reason for individual differences.

The present investigation also suggests that an observer’s belief about the world’s sequential structure is carried over from one experimental condition to another instead of being adapted to an individual condition: the model parameters derived from the randomized condition of duration reproduction provided an excellent prediction of the experimental results of the random walk condition, even though both conditions varied exactly (and only) by their sequential structure. Thus, participants in these experiments did apparently not adapt their beliefs to the actual temporal structure of the stimuli but relied on their individual hypothesis. However, whether these beliefs can be altered, for example by feedback, or reflect intrinsic personality traits warrants further investigation. A recent study on the perception of probability emphasized that average results do not provide the full picture and that individuals deviated substantially from optimal performance, with these idiosyncratic deviations persisting over a long time (Khaw et al. 2021).

Another question is whether the two-state model can encompass the full spectrum of empirical values for central tendency and sequential dependence. The two-state model predicts that sequential dependence should, with randomized stimuli, not exceed the value predicted by the simple iterative model, that is, the quadratic relationship with central tendency (red curve in Figs. 2C and 3), which has a maximum at 0.25. Indeed, this is the case for the three experiments we validated here. Note, however, that this is not a trivial result: for example, a model proposed previously to explain sequential dependence in visual orientation reproduction (Cicchini et al. 2018) predicts sequential dependence that is approximately equal to central tendency and can assume values as large as 0.5 (see SI Appendix D). While their model cannot explain the data presented here, it shows that there are alternatives to the two-state model, which would allow sequential dependence larger than 0.25. However, in our experiments, sequential dependence did not, for any of the tested participants, exceed the theoretical maximum postulated by the two-state model across three different tasks, which again suggests that our model provides a good explanation for the participants’ behaviour.

One might wonder about the purpose of integrating immediate prior information into a current decision, given that it may cause an estimation bias. One common explanation is that the regularity of our environment is relatively stable, so that integrating prior knowledge will boost the reliability of the estimation and facilitate performance (Petzschner et al. 2015, Shi et al. 2013). For a visual orientation reproduction task (Cicchini et al. 2018), the authors argued that sequential dependence provides a behavioural advantage manifesting with low reaction times and high accuracy. When the stimuli are similar between trials, it is useful to use the last perceived stimulus as prior. This assumption about the sequential structure is included in the generative assumption of the two-state model: the stimulus of the current trial is assumed to be similar to that of the last trial, since it comes from a distribution with a similar mean. However, the mean of the sampled stimuli also fluctuates over time, which makes the two-state model more flexible than a static model. That is, observers do not assume that the randomness of the external environment is strictly stable, but rather expect variations and changes.

Next, the question arises whether the proposed two-state model is optimal for the usual experimental situations with standard randomization. That is, stimuli are randomly generated as i.i.d. process from a fixed, pre-defined distribution, which has become a ‘standard’ experimental procedure since Vierordt’s work in 1868. The answer is obvious: the two-state model is not optimal, given that the stimuli are randomly drawn from a fixed distribution. Using the last trial to estimate the current would deteriorate rather than improve the quality of the estimate. However, as evidenced by the significant sequential dependence, instead of believing the stimuli are randomly generated, most of our participants assumed that there is at least some temporal continuity in the stimulus sequence. According to both the simple iterative model and the two-state model, for these participants, the overall central tendency bias should be smaller, if the stimulus sequence is changed so that stimuli are indeed similar from trial to trial. This was validated by showing in our previous study (Glasauer & Shi 2021) that the central tendency in sequences with complete random stimulus order was larger than in sequences with random-walk fluctuation. Here we showed that this decrease in central tendency and, more importantly, the remaining central tendency, is well-predicted by the two-state model on an individual basis. The model also predicts the experimentally found reversal of sequential dependence (compare the positive dependence in Fig. 5B and the negative dependence in 9B). Consequently, our model simulations together with the experimental data show that the individual assumptions about stimulus generation are stable over experimental conditions and are not adapted to the true temporal continuity (or transition probability) between stimuli.

Finally, our results show that the individual differences that are expressed in different values of central tendency and sequential dependence are not only due to differences in sensory noise, but reflect major differences in the underlying generative model, that is, in the assumptions about how stimuli are generated in the world. While some participants behave as if stimuli are generated almost independently from each other, just like when sampling from a random distribution, others show strong sequential dependence and thus assume that subsequent stimuli are similar in magnitude and depend on each other. While, on average, the perceptual system of participants seems to be optimized for random stimuli the distribution of which slowly changes over the time, the individual differences in belief about stimulus generation are not negligible.

In summary, our two-state iterative model assumes that the magnitude percept is an integration of sensory input and an updating prior knowledge. This updating can be conceived as assuming that stimuli come from a distribution the mean of which fluctuates from trial to trial. The model explains the individually different link between sequential dependence and central tendency as resulting from distinctive assumptions about the sequence structure, which differ among participants. It thus allows not only modelling the average responses of participants but also elucidates the reason for their variability: the assumptions behind the perceptual estimation process vary from person to person. The same world looks different for each of us, even when considering such a basic ability as perceiving magnitudes.

## Materials and Methods

### Duration reproduction

14 naïve volunteers (7 female, 7 male, average age 27.4) participated in the experiment, which was approved by the ethics committee of the Department of Psychology at LMU Munich. A yellow disk (diameter 4.7°, 21.7 cd/m^2^) was presented as visual stimulus on a 21-inch monitor (100 Hz refresh rate) at 62 cm viewing distance using the Psychtoolbox (http://psychtoolbox.org). Each trial started after 500 ms presentation of a fixation cross followed by the stimulus which appeared for a pre-defined duration. After a short break of 500 ms participants were prompted to reproduce the duration of the stimulus by pressing and holding a key. The visual stimulus was shown again during key press. At the end of the trial, a coarse visual feedback was given for 500 ms (5 categories from < −30% to > 30% error). Each participant performed two blocked sessions in balanced order. In the random walk condition, participants received 400 stimuli generated by cumulative summation (integration) of randomly distributed values from a normal distribution with zero mean and a SD that was chosen to yield stimuli between 400 ms and 1900 ms. In the randomized condition, the same 400 stimuli were used in scrambled order. Each participant received a different sequence (see Fig. 4A for an example). The data have been used previously (Glasauer & Shi 2021) and are publicly available (Glasauer & Shi 2021b).

### Distance reproduction

The experimental procedure has been published previously (Petzschner & Glasauer 2011) and the data are publicly accessible (Petzschner & Glasauer 2020). Briefly, 14 volunteers (7 female, 7 male, aged 22–34 years) participated. Stimuli were presented in darkness on a computer monitor as real-time virtual reality using Vizard 3.0 (Worldviz) depicting an artificial stone desert consisting of a textured ground plane, 200 scattered stones placed randomly, and a textured sky. Participants used a joystick to navigate. Estimation of travelled distances and of turning angles was tested separately under three different conditions (different ranges of distances or angles, see Fig. S5 and S6, 200 trials per condition) in a production-reproduction task. For distance estimation, participants were instructed to move forward on a linear path until movement was stopped when reaching the randomly selected production distance (same sequence for all subjects) and then had to reproduce the perceived distance in the same direction using the joystick and indicate their final position via button press. Velocity was kept constant during movement but randomized up to up to 60% to exclude time estimation strategies. No feedback was given. For angular turning estimation, the procedure was the same except that subjects had to turn.

### Data analysis: central tendency and sequential dependence

To quantify central tendency, a linear least-squares regression was fitted to stimulus reproduction plotted over stimulus duration for each participant individually using Matlab (The Mathworks, Natick MA, USA). Central tendency was defined as 1-slope of the regression line. Sequential dependence was assessed by fitting a linear least-squares regression to the error in trial *k* plotted over the stimulus in trial *k*-1 (Holland and Lockhead 1968). Note that in the literature the sequential dependence (also called serial dependence) is often quantified as current error plotted over the difference between previous and current stimulus (e.g., Fischer & Whitney 2014, Bliss et al. 2017, Kiyonaga et al. 2017, Clifford et al. 2018, Cicchini et al. 2018). However, this method is not appropriate for stimuli on an open linear scale as in the present work (see SI Appendix A).

### Modelling: Static Bayesian model

Given a set of stimuli *x_i_* drawn from a normal distribution on an open scale with mean 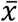, a simple *static* model for the perceptual response *y_i_* would be:

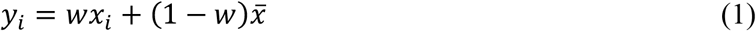

with the weight *w* being determined by using the variance of the stimulus distribution and the variance of the measurement noise. Note that the model assumes that *y_i_* only depends on the current stimulus *x_i_*, but not on the previous one. The fixed prior of the model could be the mean of the stimulus distribution 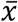. In this model, the central tendency is given as *c* = 1 − *w*. Since in this model the current response does not depend on the previous stimulus, the sequential dependence is zero regardless of the central tendency (see SI Appendix A2).

### Modelling: Iterative Bayesian model

For an *iterative* or *dynamic* model, the quantification of sequential dependence should yield an effect, given that in such a model the actual response is defined to depend on both the current and the previous magnitudes. The simplest iterative Bayesian model (Fig. 1) can be derived from two assumptions for the underlying generative process (Glasauer 2019): 1) the stimulus at the current trial is the same as the one on the previous trial plus some random fluctuation and the sensation of the stimulus is corrupted by measurement noise. For normally distributed fluctuations and noise, the Bayesian optimal estimator model can be written as Kalman filter. When the Kalman gain *k* of the model reaches its steady state (usually after few trials), its equations simplify to a weighted average, so that the response *y_i_* at trial *i* becomes

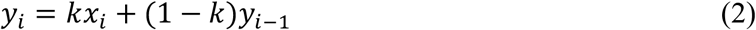

with *x_i_* being the current measurement of the stimulus and *y*_*i* − 1_ the estimate at the preceding trial *i* − 1 (Glasauer 2019). Note that for a fixed *k* this model is equivalent to the so-called “internal reference model” (Dyjas et al. 2012, Bausenhart et al. 2014).

For this iterative model, the relationship between the central tendency and the sequential dependence *s* can be determined analytically for randomly presented stimuli (see SI Appendix A3) as

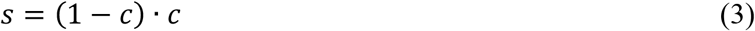

According to Eqn. 2, the central tendency is given as *c* = 1 − *k*. Intuitively, the extreme case with *k* = 0 causes the current response to completely depend on the initial response *y*_0_ (which may be arbitrary) and does not change at all; therefore, the sequential dependence becomes zero. On the other extreme, with *k* = 1 the response is veridical (*y_i_* = *x_i_*), always equal to the current stimulus, and independent of the previous, which also yields zero sequential dependence. The maximum expected sequential dependence is 0.25 for central tendency 0.5 (see Fig. 2C, orange curve). Thus, for central tendencies found experimentally, there exists a distinct testable difference between the static model (sequential dependence 0 and independent of central tendency) and the simple iterative model.

### Generative assumptions

Here we reconsider the difference between the generative assumptions of the static and iterative models. In both models, measurement noise *r* corrupts the actual sensory input. Thus, it is helpful to estimate the stimulus using additional prior information.

- The *static* model assumes that the stimulus *x_i_* in trial *i* comes from a distribution *D*(*m, v*) with a constant mean *m* and variance *v*. We thus can write the generative model as *x_i_* = *m* + *ε_x_* with *ε_x_* being a random number coming from a distribution *D*(0, *v*).
- The *iterative* model assumes that the stimulus *x_i_* in trial *i* is the same as in trial *i* − 1 except for some random change with variance *q*. In other words, the generative model is *x_i_* = *x*_*i* − 1_ + *ε_m_* with *ε_m_* coming from a distribution *D*(0, *q*).

From these assumptions we can construct a third generative model, the two-state model, that combines advantages of both models:

- The *two-state* model assumes that the stimulus *x_i_* in trial *i* comes from a random distribution *D*(*m*_*i* − 1_, *v*) with mean *m*_*i* − 1_ and variance *v*. The mean of this distribution in trial *i* is the same as in trial *i* − 1 except for some random change with variance *q*. In other words, the stimulus distribution in the current trial depends on that in the previous trial. The generative model now has two states: the randomly changing mean of the stimulus distribution *m_i_* = *m*_*i* − 1_ + *ε_m_* and the actual stimulus *x_i_* = *m*_*i* − 1_ + *ε_m_*, drawn from this distribution.

For an illustration of the generative models see SI Appendix C.

### Modelling: The two-state model

Thus, the generative equations for the two-state model are given as follows:

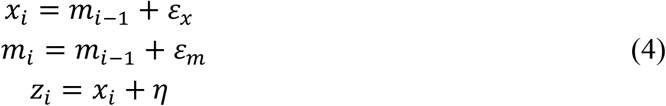

with *x_i_* being the stimulus at trial *i* that is drawn from a distribution with mean *m*_*i* − 1_ and variance *v* (here expressed by the random number *ε_x_*, which is normally distributed as *N*(0, *v*)). The mean of this stimulus distribution *m_i_* at the trial *i* is the same as in the trial before except for the random fluctuation *ε_m_* (*ε_m_* is normally distributed as *N*(0, *q*)). The actual sensory measurement (or sensation) *z_i_* is the stimulus corrupted by the sensory noise *η*, which is normally distributed as *N*(0, *r*).

We can rewrite these equations in matrix notation with 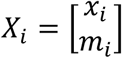 and 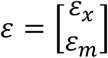, so that

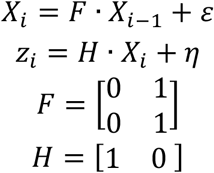

The optimal estimator for this model can be written as time-discrete Kalman filter with the covariance matrix 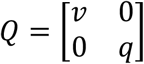 and noise variance *r*:

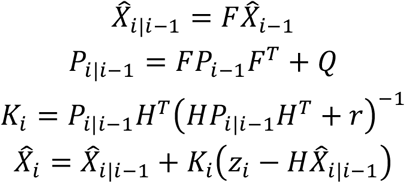

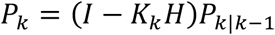

The steady state with constant matrix K thus becomes

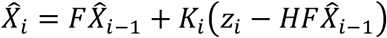

Written as states *x* and *m*, this can be expressed as

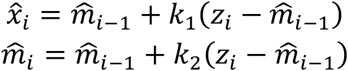

Free parameters of the model so far are the variance ratios *v*/*r* and *q*/*r*.

It should be noted that the model operates in log space (Petzschner & Glasauer 2011; Roach et al. 2017) to account for the Weber-Fechner law. Raw sensory input *d_i_* is thus transformed logarithmically to yield *z_i_* with 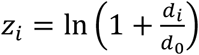. The stimulus estimate 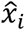 is finally back-transformed to yield 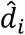 with 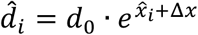. The shift term Δ*x* accounts for possible choices of the cost function and is the third free parameter of the full model.

To summarize, the two-state model has three free parameters:

1. the ratio of the variances v and r indicating the variability of the stimulus distribution relative to the sensory noise,
2. the ratio of variances q and r indicating the variability of the additive random shift relative to the sensory noise, and
3. a shift parameter that accounts for global over- or underestimation (see also Petzschner & Glasauer 2011).

For all three models (static, simple iterative, two-state) the same model equations and the same Kalman filter can be applied. The three models differ by the free parameters:

1. static model: *ε_m_* = 0, therefore variability *q* = 0. Free parameters: *v*/*r* and Δ*x*.
2. iterative model: *ε_x_* = 0, therefore variability *v* = 0. Free parameters: *q*/*r* and Δ*x*.
3. two-state model: full model. Free parameters: *q*/*r*, *v*/*r*, and Δ*x*.

### Model fitting and model selection

For model simulation, the individual stimulus sequences were used to fit the model separately for each participant. Thus, the model received the sequence of stimuli in exactly the same order as the participant and computed a sequence of responses. Model fitting was performed in linear stimulus space, that is, for model fitting, the least-squares distance between stimulus sequence and responses was minimized. The Matlab function *lsqnonlin* was used to estimate the parameters, and *nlparci* was applied to estimated confidence intervals. The coefficient of determination R^2^ for average data was calculated as 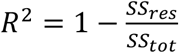, with SS_res_ being the residual sum-of-squares and SS_tot_ the total sum-of-squares. If a model perfectly captures the data, *R*^2^ = 1. Models with negative *R^2^* are worse than the baseline model, which predicts the average of the data and which has *R*^2^ = 0.

To compare models, we used a leave-one-out (LOO) cross-validation procedure (Arlot & Celisse 2010) adapted to time series. In LOO, each of the n data points is successively “left 20 out” from the sample and used for validation by fitting the model to the remaining data points and recording the error of the left-out data point. The criterion is the average validation error of the n model fits: the best model is the one with the minimal validation error. To account for the trial-to-trial dependence of data or model, in the modified LOO instead of leaving out just one data point, k values around this data point are left out, while the validation error is only computed for the data point in the center of this leave-out window. Here we selected k=11, which was assumed to be large enough to account for trial-to-trial dependencies. The same result (8x two-state model selected) was already achieved with k=3, while for k=1 the two-state model was best in 9 cases.

## Acknowledgements

This work was supported by German Research Foundation (DFG) grants GL 342/3-2 and SH 166/3-2. We thank Mauro Manassi, David Whitney, and Jason Fischer for pointing out that for stimuli on a circular scale the RSD is valid without causing artifacts, and that for the RSD permutation tests should be run as statistical sanity check.

## Data availability statement

Data used in this publication are freely available as Glasauer & Shi (2021b) https://doi.org/10.12751/g-node.hdsam3 and Petzschner & Glasauer (2020), https://doi.org/10.12751/g-node.21796b.

## Author contributions

SG and ZS wrote the main manuscript text and prepared the figures. All authors reviewed the manuscript.

## Footnotes

1 In the Bayesian estimation process, the estimate of the current stimulus is based on a posterior distribution, which is the product between the likelihood distribution (describing the sensory uncertainty that is inherited from the current sensory input) and the prior distribution (describing prior knowledge). The estimate can be the most likely value from the posterior distribution, the maximum of the distribution, or other measures such as the mean, and its uncertainty can also be derived.

2 Model simulation performed after transforming the stimulus data to the log-domain, as done previously (Petzschner & Glasauer 2011; Roach et al. 2017), to account for the dependence of the variance on stimulus size (Weber-Fechner law). Fitting was done by minimizing the least-squares distance between trial-wise response and simulation in the linear domain.

