## Supplemental material for "Individual beliefs about temporal continuity explain variation of perceptual biases"

### Appendix A. The relative and absolute sequential dependence and their relations to the Bayesian models.

In the literature, two main methods to quantify sequential (or serial) dependence can be found. Older studies such as Holland and Lockhead (1968) quantified the sequential effect as the dependence of the current error on the stimulus magnitude in the previous trial (we refer to it as the absolute sequential dependence, ASD). More recent studies (e.g., Fischer & Whitney 2014, Bliss et al. 2017, Kiyonaga et al. 2017, Clifford et al. 2018, Cicchini et al. 2018) reported the dependence of the current error on the difference between the stimuli in the previous and current trials (the relative sequential dependence, RSD).

Even though it is rarely mentioned, the RSD is appropriate only when stimuli 1) come from a circular scale, such as angular orientation, and 2) are equally distributed over the whole scale. This is the case for most of the studies mentioned above, which investigated serial dependence for perception of visual orientation. For other cases, such as when stimuli are drawn from an open scale or from only part of a circular scale, the RSD is problematic and potentially misleading, because it inflates the true sequential dependence effect (see details in Appendix A2 below). Moreover, the RSD can then falsely show a dependence, based on mathematical coupling (Archie 1981, Curran-Everett 2010), even if the result of the current trial is completely independent of the previous stimulus.

#### *A1. Relative and absolute serial dependence for a special case*

To show that relative sequential dependence (RSD) is not appropriate for stimuli on an open scale, such as distance or duration, let us consider a typical perceptual experiment (Fig. S1A): stimuli are presented successively in randomized order, and for each stimulus one response is measured, that should ideally reflect the stimulus magnitude (typical responses shown in Fig. S1B). Now consider a 'dead salmon case' (a famous fMRI study showing BOLD responses in a dead salmon, Bennett et al. 2009): if the response is constant, say zero, independently of the given stimuli (Fig. S1C), we still get a strong 'serial dependence' in terms of RSD measure (Fig. S1D). The RSD also remains strong when the response is random with no relation to the stimulus (Fig. S1E).

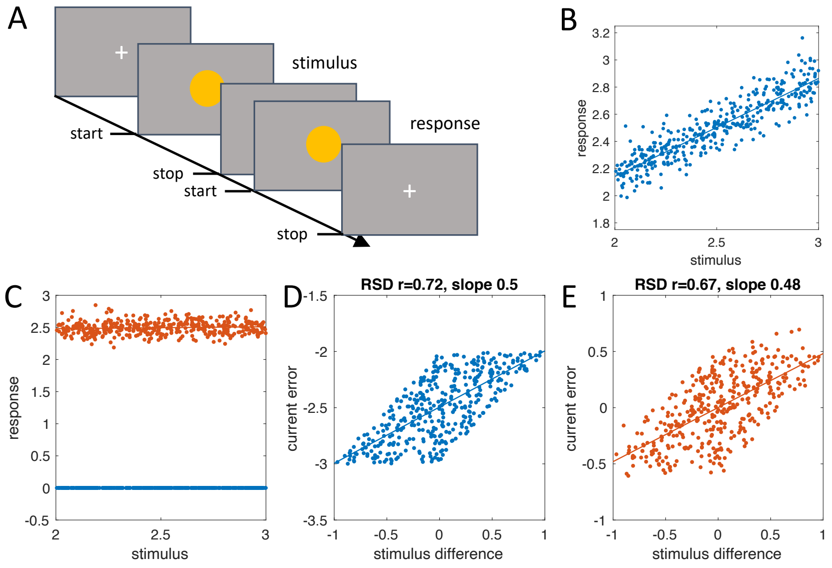


**Figure S1**: Simulations of a perceptual experiment. **A**: Schematic timeline of a typical duration production-reproduction experiment. Stimulus duration given by a visual cue is actively reproduced as response. **B**: Exemplary responses plotted over stimulus for 400 randomly presented stimuli with an arbitrary stimulus range of 2-3. **C**: "dead salmon" case (blue, zero response) and independent random responses with a central tendency of 1 (red dots) to the same stimuli as in A. **D**: the RSD measure for the zero response in C shows a highly significant relation with a correlation coefficient of r=0.71 and a regression slope of 0.5, erroneously suggesting a serial dependence. **E**: again, the corresponding RSD shows a significant relation. Note, however, that an appropriate statistical permutation test reveals for both cases that both slope and correlation coefficient are not statistically significant (here: both p>0.3).

Why is that the case? Because the current stimulus magnitude is calculated twice in both the response error (on the y-axis) and the stimulus difference (on the x-axis). Thus, the two variables are algebraically coupled, which engenders a strong bogus correlation between both variables. For the 'dead salmon' case in Fig. S1C, it can be shown analytically that the slope of the regression line of the RSD becomes 0.5.

Let us denote $y_i$ as the current response and $x_i$ as the current stimulus of the trial and $i$. For the "dead salmon" case, response is "dead" ($y_i = 0$) and the error plotted on the y-axis is $e_i = y_i - x_i = -x_i$. The inter-trial difference plotted on the x-axis is $d_i = x_{i-1} - x_i$ (as defined in Fischer & Whitney 2014). The regression slope of the two variables is calculated by $m = \frac{\sum_{i=2}^{n}(d_i-\bar{d})(e_i-\bar{e})}{\sum_{i=2}^{n}(d_i-\bar{d})^2}$. Since the stimuli $x_i$ are presented in random order, the mean difference $\bar{d}$ approaches zero for large n. Consequently, $m = \frac{\sum_{i=2}^{n}(d_i)(-x_i+\bar{x})}{\sum_{i=2}^{n}(d_i)^2} = \frac{\sum_{i=2}^{n}(x_{i-1}-x_i)(-x_i+\bar{x})}{\sum_{i=2}^{n}(x_{i-1}-x_i)^2}$. The denominator can be rewritten as

$$\sum_{i=2}^{n}(x_{i-1}-x_i)^2 = \sum_{i=2}^{n}(x_{i-1}^2 - 2x_i x_{i-1} + x_i^2) \approx 2\sum_{i=2}^{n}(x_i^2 - x_i x_{i-1})$$

and the numerator becomes

$$\sum_{i=2}^{n}(x_{i-1}-x_i)\,(-x_i+\bar{x}) = \sum_{i=2}^{n}(x_i^2 - x_{i-1}x_i) + \bar{x}\sum_{i=2}^{n}(x_{i-1}-x_i) \approx \sum_{i=2}^{n}(x_i^2 - x_i x_{i-1})$$

Consequently, the slope becomes $m = \frac{1}{2}$ for large enough n.

Note that this holds for random stimuli on an open scale, but not on a circular scale as used in Fischer & Whitney (2014) and many other recent studies on serial dependence. For the circular scale, if stimuli are drawn from the whole circular range, the slope becomes zero even for the "dead salmon" case.

*A2. Relative and absolute serial dependence for a Bayesian model with fixed prior*

Let's consider a Bayesian model with a fixed static prior for the central tendency which should not show a serial dependence, because the estimate in the current trial $y_i$ is, in first approximation, generated by a weighted average of the measurement of the current stimulus $x_i$ and a fixed value (usually the mean $\bar{x}$ of the stimuli, assuming a normal or log-normal distribution of the stimuli). The current response can then be computed by

$$y_i = wx_i + (1 - w)\bar{x}$$

with $w$ being a weight between 0 and 1. The model predicts a central tendency $c = 1 - w$ with large magnitudes being underestimated and small ones overestimated, no matter what the previous stimulus is. The range of the stimuli, however, causes a sequential structure of the trials with a high probability of a large magnitude followed by a small magnitude, and vice versa. Thus, the RSD, measured by the inter-trial difference, *falsely* takes the central tendency effect as the sequential dependence. Fig S2A shows a simulation of the static model with a noisy response that yields a more realistic dependence of the result on the stimulus. Again, the simulation of the RSD shows a significant relationship (Fig. S2B) despite the error being independent of the previous stimulus for the static model (Fig. S2C). Evidently, the RSD measure used in the recent literature could be dangerously misleading for such a case on an open scale. In the presence of a central tendency (or other systematic errors), the RSD measure without appropriate permutation testing cannot make clear assertions on whether the response in the actual trial really depends on the stimulus in the previous trial or not.

We show in the following that the RSD in a model with fixed prior results in a dependence of error on stimulus difference that could be mistaken as serial dependence. Given a set of stimuli $x_i$ drawn from a random distribution on an open scale with mean $\bar{x}$, a simple model for the perceptual response $y_i$ would be:

$$y_i = wx_i + (1 - w)\bar{x}$$

with the weight $w$ being determined, for example, by using the variance of the stimuli and the variance of the measurement noise. Note that the model assumes that $y_i$ only depends on the current stimulus $x_i$, but not on the previous one. In this model, the central tendency is given as $c = 1 - w$. The RSD (serial dependence according to Fischer & Whitney 2014) is a systematic dependence of the error $x_i - y_i$ on the stimulus difference $x_{i-1} - x_i$. Note that in Fischer & Whitney (2014) a derivative-of-Gaussian was fitted to the orientation judgements as orientation difference has lower and upper bounds. We show here for unbound continuous magnitude using a linear fit that with this formulation a systematic dependence arises even for the trial-independent model above.

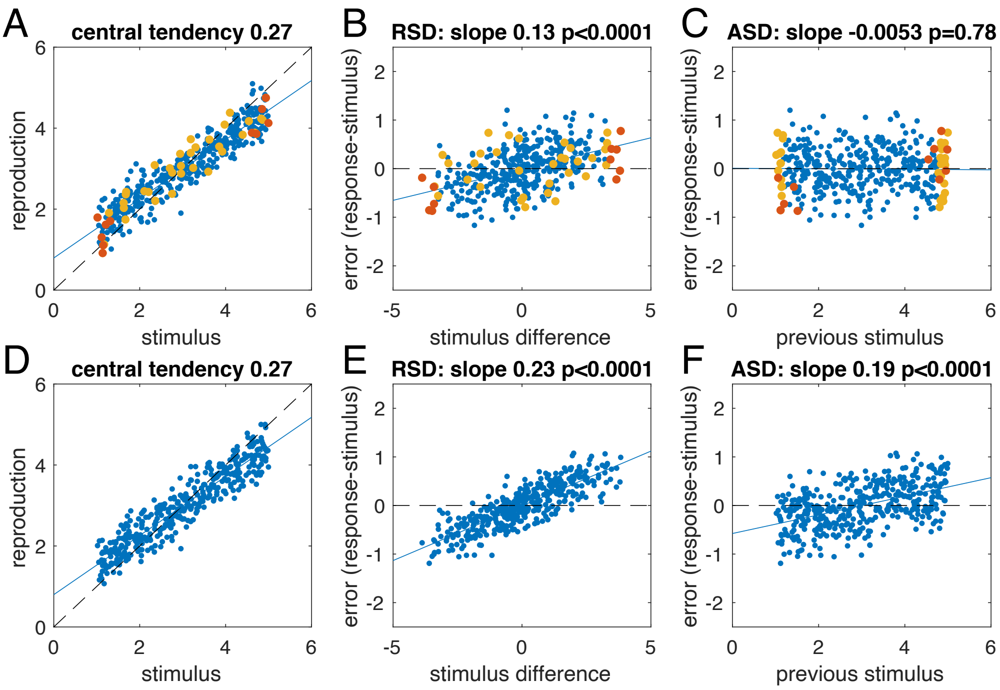


**Figure S2**: Simulation of two Bayesian models estimating a random sequence of 400 stimuli presented in the magnitude range 1-5. (**A**-**C**) Simulation of a Bayesian model with fixed prior. The actual stimuli have been corrupted with normally distributed measurement noise mimicking random fluctuations in sensation. **A**: reproduced magnitude plotted over stimulus; the central tendency is given as 1-slope of the regression line (blue). **B**: Relative serial dependence (RSD): Error (response-stimulus) in the current trial plotted over the difference between stimuli in the previous and the current trials to quantify serial dependence (the misleading p-value is calculated from standard regression statistics). **C**: Absolute serial dependence (ASD): Error (response-stimulus) plotted over stimulus in the previous trial. The serial dependence, given as slope of the regression, now correctly reports a value that is not significantly different from zero, as expected for a model with a fixed prior. Simulated data points with large inter-trial stimulus differences (red, on right and left sides in B) or previous stimulus with extreme magnitudes (yellow, on the right and left sides in C) are marked in colour to emphasize that large errors correlate with large stimulus differences. Even though no true serial dependence exists in this simulation, the RSD slope differs significantly from zero and amounts to approximately 50% of the central tendency. (**D**-**F**) Simulation of an iterative Bayesian model estimating the same random sequence of 400 stimuli as in Fig. S2A-C and yielding the same central tendency. D: reproduced magnitude plotted over stimulus; the central tendency is given as 1-slope of the regression line (blue). E: Relative serial dependence (RSD): Error (response-stimulus) in the current trial plotted over the difference between stimuli in the previous and current trials. Note that the slope is larger than for the static model (Fig. 2B) F: Absolute serial dependence (ASD). Error (response-stimulus) plotted over stimulus in the previous trial. The serial dependence, given as slope of the regression, differs significantly from zero, as expected from the iterative model.

The linear relation is given by $y'_i = mx'_i + b$ or $y_i - x_i = m(x_{i-1} - x_i) + b$. Thus, the slope of the best fit using least-squares is $m = \frac{\sum_{i=2}^{n}(x'_i - \overline{x'})(y'_i - \overline{y'})}{\sum_{i=2}^{n}(x'_i - \overline{x'})^2}$. Since the stimuli are randomly distributed, $\overline{x'} = \frac{1}{n}\sum_{i=2}^{n}(x_{i-1} - x_i) \approx 0$ for large enough sample size n.

Similarly, $\overline{y'} = \frac{1}{n}\sum_{i=2}^{n}(y_i - x_i) = \bar{y} - \bar{x} \approx 0$, because $\bar{y} = \frac{1}{n}\sum_{i=1}^{n}(wx_i + (1-w)\bar{x}) = \bar{x}$.

With the equation of the static model, the slope can be rewritten as

$$m = \frac{\sum_{i=2}^{n} x'_i y'_i}{\sum_{i=2}^{n} x'^2} = \frac{(1-w)\sum_{i=2}^{n}(x_i - x_{i-1})(x_i - \bar{x})}{\sum_{i=2}^{n}(x_i - x_{i-1})^2} = (1-w)\frac{\sum_{i=2}^{n}(x_i - x_{i-1})x_i}{\sum_{i=2}^{n}(x_i - x_{i-1})^2}$$

Note that the denominator can be rewritten as

$$\sum_{i=2}^{n}(x_i - x_{i-1})^2 = \sum_{i=2}^{n}(x_i{}^2 - 2x_i x_{i-1} + x_{i-1}{}^2)$$

$$\approx 2\left(\sum_{i=2}^{n} x_i{}^2 - \sum_{i=2}^{n} x_i x_{i-1}\right) = 2\sum_{i=2}^{n}(x_i - x_{i-1})x_i$$

And therefore, we have the following relation

$$m \approx \frac{1-w}{2} = c/2$$

That is, the half of the central tendency effect is falsely attributed to the RSD, even though the model assumes no sequential dependence.

If, for the static model above, the current error $(x_i - y_i)$ is plotted over the last stimulus $x_{i-1}$, then the slope of the best fit regression line becomes $m' = \frac{\sum_{i=2}^{n}(x_{i-1}-\bar{x})(x_i-y_i)}{\sum_{i=2}^{n}(x_{i-1}-\bar{x})^2}$. While the denominator is the autocorrelation of $(x - \bar{x})$ with lag zero, the numerator can be evaluated as follows:

$$\sum_{i=2}^{n}(x_{i-1} - \bar{x})(x_i - y_i) = \sum_{i=2}^{n}(x_{i-1} - \bar{x})(x_i - wx_i - (1-w)\bar{x})$$

$$= (1-w)\sum_{i=2}^{n}(x_{i-1} - \bar{x})(x_i - \bar{x})$$

Therefore, the numerator is the autocorrelation of $(x - \bar{x})$ with lag 1 weighted by the central tendency $(1 - w)$. The slope $m'$ therefore depends on the model weight (or the central tendency) but can maximally assume a value that is the quotient between stimulus autocorrelations with lag 1 and lag 0. Since the stimulus autocorrelation for lag 1 is close to zero for random stimuli, the serial dependence is also close to zero, as expected for a model in which the current response is independent of previous stimuli.

*A3. Serial dependence for an iterative Bayesian model*

The simplest probabilistic model for serial dependence is a Kalman filter model with the assumption that the stimulus at the current trial is equal to that of the last trial plus some random fluctuation (e.g., Glasauer 2019). Since the Kalman gain *k* reaches a steady state value relatively quickly, the basic iterative equation can be simplified to

$$y_i = kx_i + (1-k)y_{i-1}$$

Fig. S2D shows the simulated response of the iterative model to the same stimuli as in Fig. S2A. For better comparison, the parameter *k* was chosen to approximate the central tendency to that of the static model. The RSD again shows a significant positive slope (Fig. 2SE), which is larger than in Fig. S2B. But now also the ASD shows a significant relationship and confirms that the current response indeed depends on the previous stimulus signifying a serial dependence.

The central tendency is given by the linear least squares fit $y_i = wx_i + b$, for which the slope *w* can be determined as $w = \frac{\sum_{i=1}^{n}(x_i-\bar{x})(y_i-\bar{y})}{\sum_{i=1}^{n}(x_i-\bar{x})^2}$. We first determine the mean of the responses as $\bar{y} = \frac{1}{n}\sum_{i=1}^{n} y_i = \frac{1}{n}\sum_{i=1}^{n}[kx_i + (1-k)y_{i-1}] = k\bar{x} + (1-k)\frac{1}{n}\sum_{i=1}^{n} y_{i-1} \approx k\bar{x} + (1-k)\bar{y}$

From this directly follows that $\bar{y} \approx \bar{x}$ .
The denominator can be rewritten as follows

$$\sum_{i=1}^{n}(x_i - \bar{x})^2 = \sum_{i=1}^{n}(x_i^2 - 2x_i\bar{x} + \bar{x}^2) = \sum_{i=1}^{n} x_i^2 - 2\bar{x}\sum_{i=1}^{n} x_i^2 + n\bar{x}^2 = \sum_{i=1}^{n} x_i^2 - n\bar{x}^2$$

The numerator can be written as

$$\sum_{i=1}^{n}(x_i - \bar{x})(y_i - \bar{y}) = \sum_{i=1}^{n}(x_i y_i - x_i\bar{y} - \bar{x}y_i + \bar{y}\bar{x}) = \sum_{i=1}^{n} x_i y_i - n\bar{y}\bar{x}$$

$$= \sum_{i=1}^{n}\left(x_i(kx_i + (1-k)y_{i-1})\right) - n\bar{y}\bar{x} = k\sum_{i=1}^{n} x_i^2 + (1-k)\sum_{i=1}^{n} x_i y_{i-1} - n\bar{y}\bar{x}$$

$$= k\sum_{i=1}^{n} x_i^2 - k\sum_{i=1}^{n} x_i y_{i-1} + \sum_{i=1}^{n} x_i y_{i-1} - n\bar{y}\bar{x}$$

$$= k\sum_{i=1}^{n} x_i^2 - k\sum_{i=1}^{n} x_i y_{i-1} = k\sum_{i=1}^{n} x_i^2 - kn\bar{y}\bar{x} = k\left(\sum_{i=1}^{n} x_i^2 - n\bar{x}^2\right)$$

because $\sum_{i=1}^{n} x_i y_{i-1} - n\bar{x}\bar{y} = 0$ since $x_i$ is independent of $y_{i-1}$ and therefore $\sum_{i=1}^{n} x_i y_{i-1} = n\bar{x}\bar{y} = n\bar{x}^2$. Thus, $w = k$ and $b = \bar{x}$. In other words, the central tendency $c = 1 - w = 1 - k$ reflects the steady state Kalman gain of this iterative model.

The slope of the error plotted over the last stimulus is $m = \frac{\sum_{i=2}^{n}(x_{i-1}-\bar{x})(x_i - y_i)}{\sum_{i=2}^{n}(x_{i-1}-\bar{x})^2}$. From the slope of the linear regression $w = \frac{\sum_{i=1}^{n}(x_i-\bar{x})(y_i-\bar{y})}{\sum_{i=1}^{n}(x_i-\bar{x})^2}$ , it follows that the denominator can be expressed as

$$\sum_{i=1}^{n}(x_i - \bar{x})^2 = \frac{1}{w}\sum_{i=1}^{n}(x_i - \bar{x})(y_i - \bar{y})$$

The numerator can be rewritten as follows:

$$\sum_{i=2}^{n}(x_{i-1} - \bar{x})(x_i - y_i) = \sum_{i=2}^{n}(x_{i-1} - \bar{x})(x_i - kx_i - (1-k)y_{i-1})$$

$$= (1-k)\sum_{i=2}^{n}(x_{i-1} - \bar{x})(x_i - y_{i-1})$$

Therefore

$$m = \frac{\sum_{i=2}^{n}(x_{i-1} - \bar{x})(x_i - y_i)}{\sum_{i=2}^{n}(x_{i-1} - \bar{x})^2} = \frac{(1-k)\sum_{i=2}^{n}(x_{i-1} - \bar{x})(x_i - y_{i-1})}{\frac{1}{w}\sum_{i=1}^{n}(x_i - \bar{x})(y_i - \bar{y})} = (1-w)w$$

This relation between serial dependence and central tendency is shown as red curve in Figs. 2 and 3.

### Appendix B. Generative models

The assumptions behind the three estimation models can be illustrated by looking at the generative mechanisms assumed by the estimation models. These mechanisms are also called generative models (e.g., Griffiths et al. 2008) and illustrated in Fig. S3. Each generative model yields different predictions concerning the sequential dependence. The two-state model assumes two hidden states (the stimulus and the mean of the stimulus distribution, Fig. S3B), which generalizes the *two-stage* model proposed previously (Petzschner & Glasauer 2011). The original *two-stage* model, which is not identical to the simple iterative model mentioned in the main text and above, assumes that the variance $v$ of the stimulus distribution is directly related to the variance of the known random fluctuations $q$ of the mean and that it could be estimated from these fluctuations. When this restriction is released, the resulting two-state model encompasses the *two-stage* model as a special case, and also both the static model and the simple iterative model as two boundary cases: The static model is a boundary case of the two-state model*, if the mean $m_i$ in the two-state model is constant instead of fluctuating (i.e., $q = 0$). When the assumed stimulus distribution has a negligible variance ($v = 0$), the two-state model reduces to the simple iterative model.

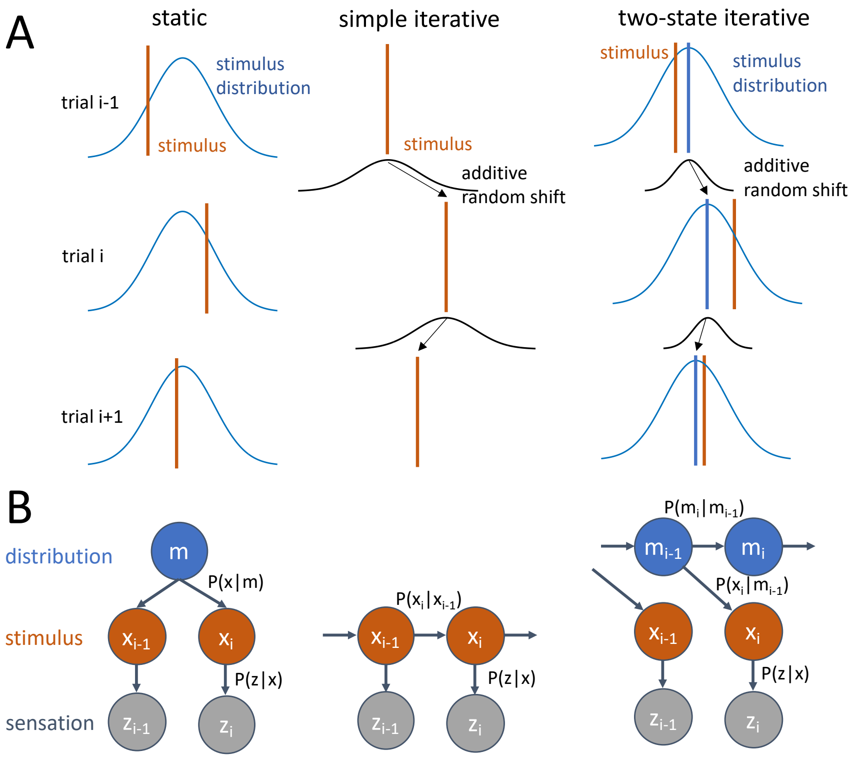


**Figure S3**: Generative models of magnitude reproduction. **A**: The static model (left column) assumes that stimuli (red lines) are randomly drawn from a fixed stimulus distribution (blue). In the simple iterative model (middle column), the stimulus at the current trial $i$ is assumed to be the same as in the previous trial $i$-1 except for some random fluctuation that is conceptualized as additive random shift (black distribution). The two-state iterative model (right column) combines these two models: the stimulus is drawn from a stimulus distribution (blue), but the mean of the stimulus distribution fluctuates from trial to trial by a random amount (additive random shift, black distribution). In the illustration the stimuli are the same in all three cases, but the assumed sequential structure

* To be exact, the two-state model with $q$=0 and the static model are equivalent only for the steady state, because the two-state model initially estimates the unknown mean of the stimulus distribution $m$, while the static model assumes that the mean $m$ is already known at trial 1.

differs in each model. The measurement noise distribution around the stimuli has been omitted for clarity. **B**: Hidden Markov Model diagram visualizing the three generative models. Note that the sensation *z*, the observable, is also shown. It is derived from the stimulus via the conditional probability $P(z|x)$, which describes the measurement noise. For the static model, the state *m*, which represents the mean of the stimulus distribution, is assumed not to change from trial to trial. Stimuli are drawn from the stimulus distribution with $P(x|m)$. In the simple iterative model, stimuli change from one trial to the next with a stationary transition probability $P(x_i|x_{i-1})$. In the two-state model, the mean of the stimulus distribution changes from trial to trial with a stationary transition probability $P(m_i|m_{i-1})$, and the stimuli are drawn from the new stimulus distribution with $P(x_i|m_{i-1})$.

### Appendix C. Experimental validation

#### *C1. Duration Reproduction*

The average data for the randomized condition of the duration reproduction experiment (14 subjects, data available as Glasauer & Shi 2021) is presented in Fig. S4. To present average data, we grouped individual data in bins of 0.1 s and then averaged over subjects. The model exhibits not only an excellent fit to the averaged data for reproduced duration plotted over stimulus duration (Fig. S4 $A_1$, coefficient of determination $R^2=.986$), but also captures the average absolute serial dependence of the error on the previous stimulus (Fig. S4 $A_2$, $R^2=.827$), even though this relation was not included in the fitting procedure.

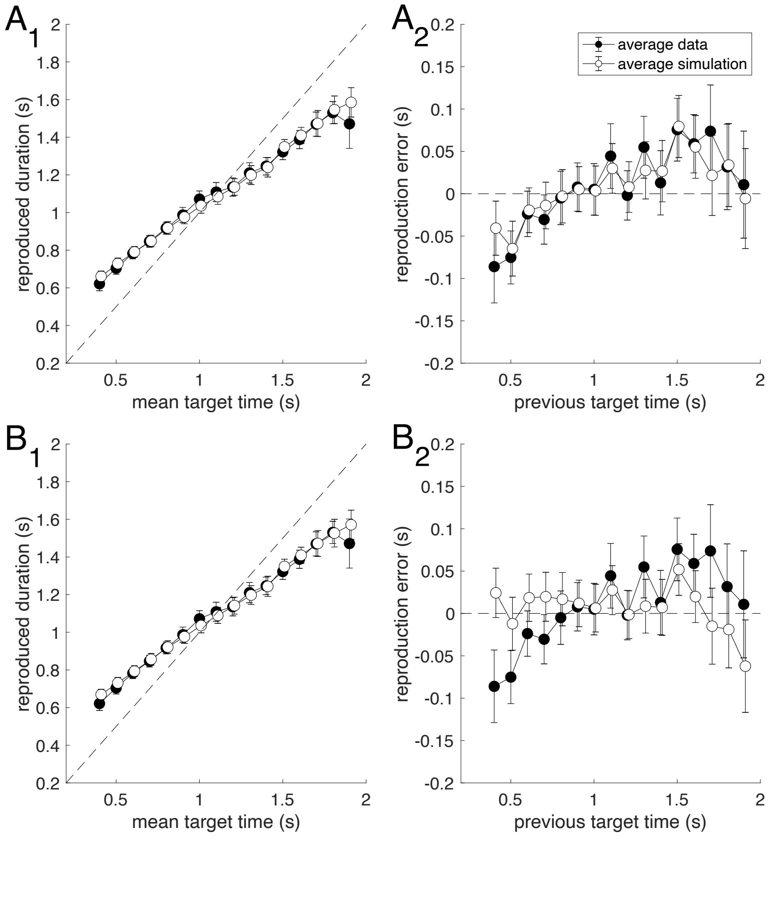

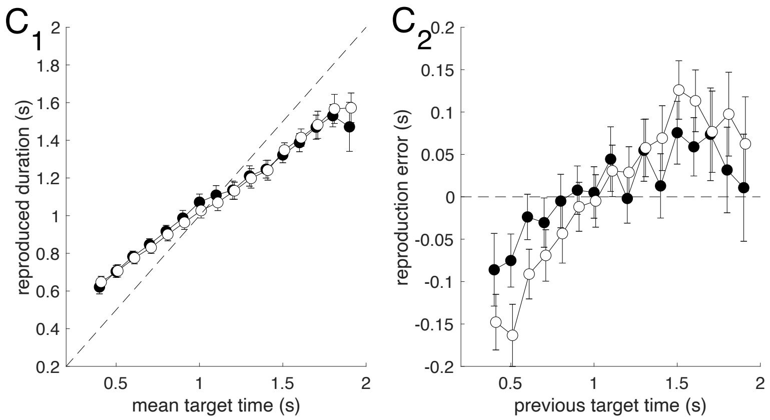


**Figure S4**: Average data for duration reproduction experiment (black dots) together with average simulation result for the best fit models (open dots). **A**: two-state model. **B**: static model. **C**: iterative model. **A1-C1**: Reproduced duration as a function of the current target duration. **A2-C2**: Reproduction error as a function of the previous target duration. Data points are averages of 7 to 14 subjects (only 7 subjects received long durations above 1.7 s); error bars denote SEM. For the model simulation, the models were fitted individually to the trial-by-trial time course of the responses of each subject. From theses individual simulated response time courses, the average simulation was computed in the same way as for the real response time courses. Note that for all models average mean data and simulated responses match well when plotted over current target time (A1-C1), but only match in A2, but not B2 and C2, when plotted over previous target time, showing that the serial dependence is not appropriately captured by the static or simple iterative models.

For comparison, the best fit simple iterative model yields $R^2=.985$ for average reproduced duration over current stimulus (Fig. S4 $C_1$), but $R^2=-.039$ for average error over previous stimulus, indicating that the model is not appropriate for explaining this relation (Fig S4 $C_2$). Similarly, the static model yields $R^2=.987$ for average reproduced duration over current stimulus (Fig. S4 $B_1$), but $R^2=-.240$ for error over previous stimulus (Fig S4 $B_2$), showing that this model is also inappropriate for explaining the serial dependence.

Fig. S5 shows the strong cross correlation in the data to the previous few trials, which is significantly different from zero up to lag 3 (t-test; lag 2: $p = 0.0007$; lag 3: $p=0.039$; n=14), and thus confirms the serial dependence in the data.

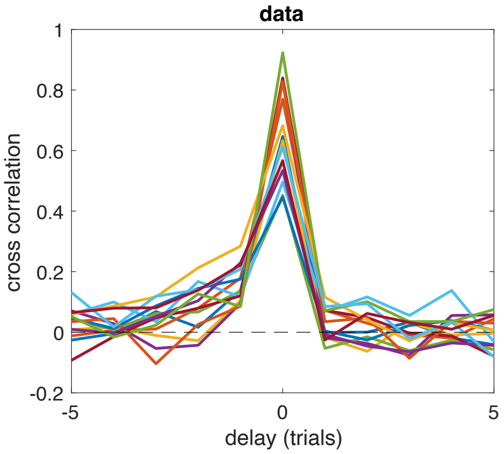


**Figure S5**: cross correlation between stimulus and reproduction for the duration reproduction experiment for all participants (n=14, colours correspond to individual participants).

The average data and model predictions for the random walk condition are shown in Fig. S6. Fig. S7 summarizes the average data for both conditions together with the model fits for the randomized condition and the model predictions for the random walk condition and includes the serial dependence up to lag 2.

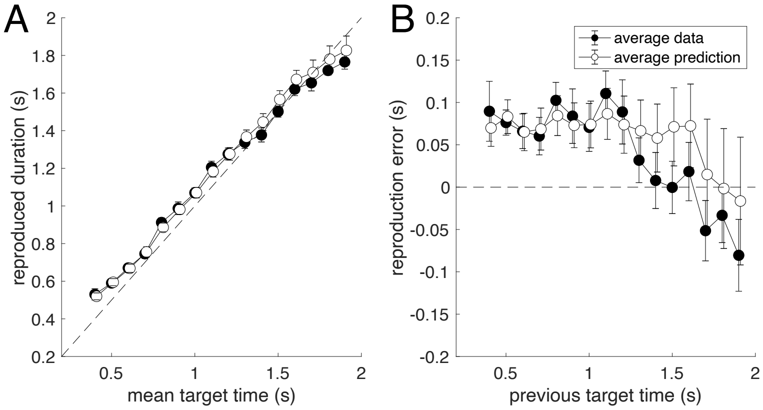


**Figure S6**: Average data for duration reproduction experiment with random-walk trial order (black dots) together with average simulation result predicted from the individual best fit iterative two-state model (open dots). **A**: Reproduced duration as a function of the current target duration. **B**: Reproduction error as a function of the previous target duration. Data points are averages of 7 to 14 subjects (only 7 subjects received long durations above 1.7 s); error bars denote SEM. For the model predictions, the individual models previously fitted to the trial-by-trial time course of the responses of each subject to the random order experiment was used without change. From the predicted individual response time courses, the average prediction was computed in the same way as for the real response time courses. Note that for this prediction the data shown here were not used, thus the good match between model prediction and data supports the validity of the model and that model parameters are preserved despite the change in stimulus sequence.

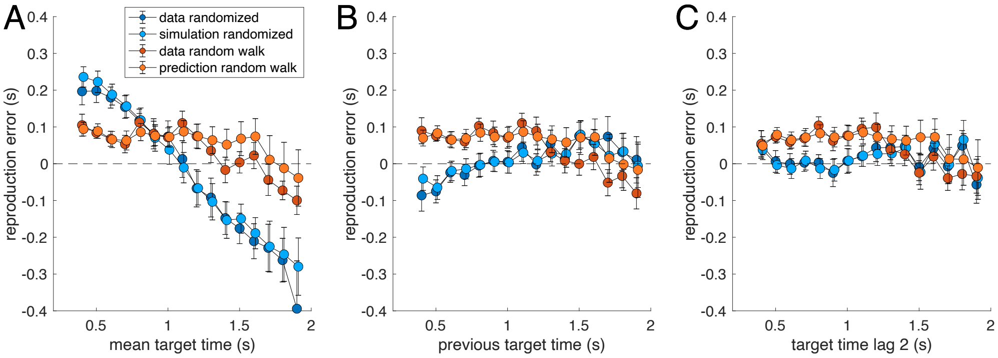


**Figure S7**: Average reproduction error for both conditions of the duration reproduction experiment (randomized: dark blue; random walk: dark orange) together with average simulation result fitted to the randomized condition (light blue) and predicted from for the random walk data (light orange). **A**: Reproduction error as a function of the current target duration. **B**: Reproduction error as a function of the previous target duration. **C**: Reproduction error as a function of the target duration two trials in the past. Error bars denote SEM. See also Fig. S6.

*C2. Distance Reproduction*

The data were published as Petzschner & Glasauer (2011) and are available as Petzschner & Glasauer (2020). Figs S8 and S9 show the average simulation of the distance and angular reproduction from the two-state model. Similar to the duration reproduction study, the two-state model also fits very well both for the central tendency and the sequential dependence.

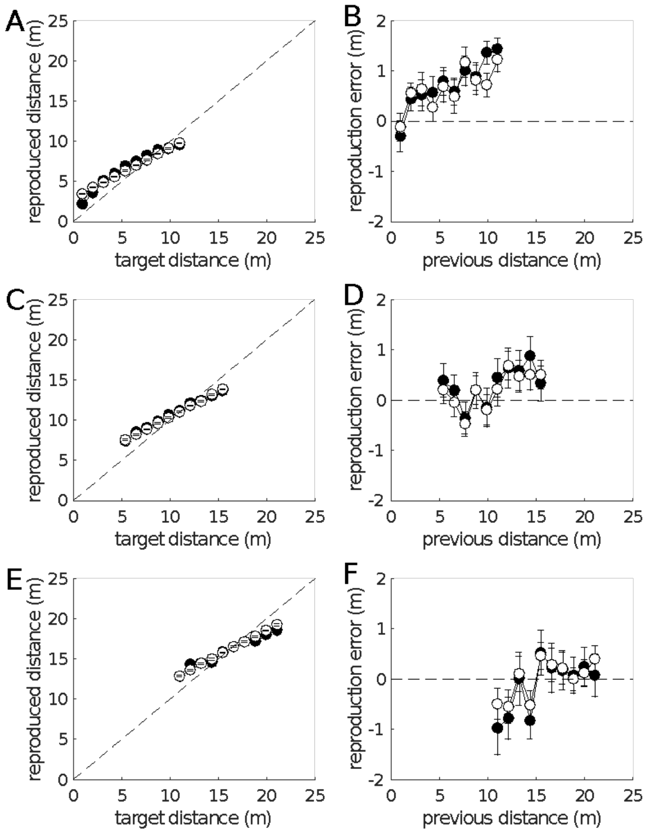


**Figure S8**: Average data for linear distance reproduction experiment (black dots, Petzschner & Glasauer 2020) together with average simulation result of the best fit iterative two-state model (open dots). Each error bar denotes a standard error of the mean. **A, B**: small range; **C, D**: middle range; **E, F**: large range. The model (three free parameters) was fit individually to the trial-by-trial time course of all responses of each subject. The three ranges are shown in separate sub-plots only for convenience. Notably, fitting a single model (three free parameters) to the average trial-by-trial time course (each subject in this experiment received the stimuli in the same order; all three ranges were presented successively) yields almost indistinguishable results (not shown).

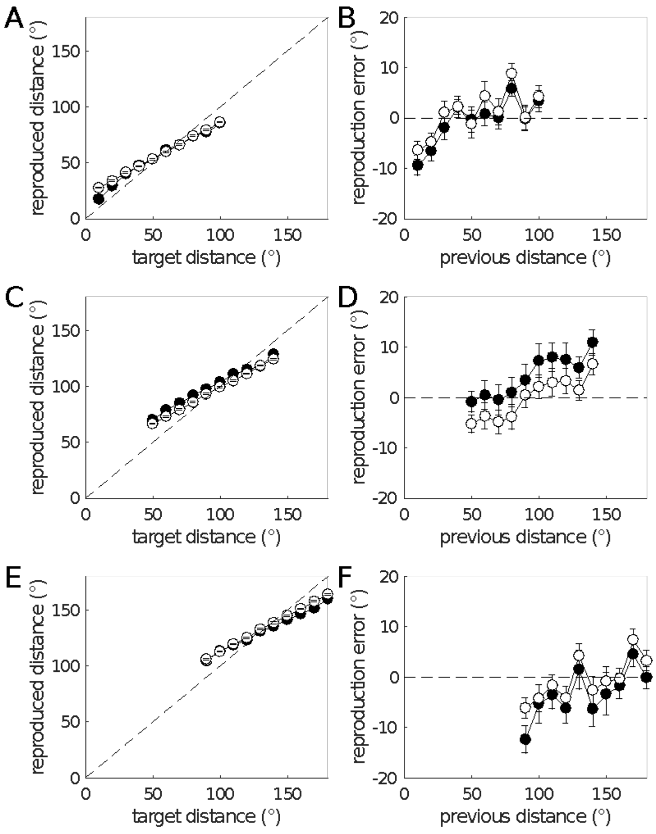


**Figure S9**: Average data for angular distance reproduction experiment (black dots) together with average simulation results of the best fit iterative two-state model (open dots). See Fig. S8 for more information.

### Appendix D. The model of sequential dependence by Cicchini et al. (2018)

Another model for serial dependence has been proposed in Cicchini et al. 2018 for orientation perception on a circular scale (as in Fischer & Whitney 2014). Here we test whether their model can explain the serial dependence seen in our example data for magnitude perception on linear scales.

In its basic form, without considering that the weight $w$ itself is considered to be dependent on the "distance between the two cues" $d$, the model can be expressed as

$$y_i = wx_i + (1 - w)x_{i-1}$$

The central tendency of this model is given by $c = 1 - w$, because $\bar{y} = \bar{x}$ for a random stimulus sequence. Reformulating the model equation yields $y_i - x_i = (1 - w)(x_{i-1} - x_i)$. Thus, the RSD slope of this model is equal to the central tendency. It also follows that the correct quantification of the serial dependence by the ASD yields the same slope $(1 - w)$, because the least squares fit $y_i - x_i = mx_{i-1} + b$ yields $m = 1 - w$ (Fig. S10).
Note that despite the apparent similarity of this model to the optimal Bayesian estimator (simple iterative model, see above), the two models are very different in their predictions for serial dependence and its relation to central tendency.

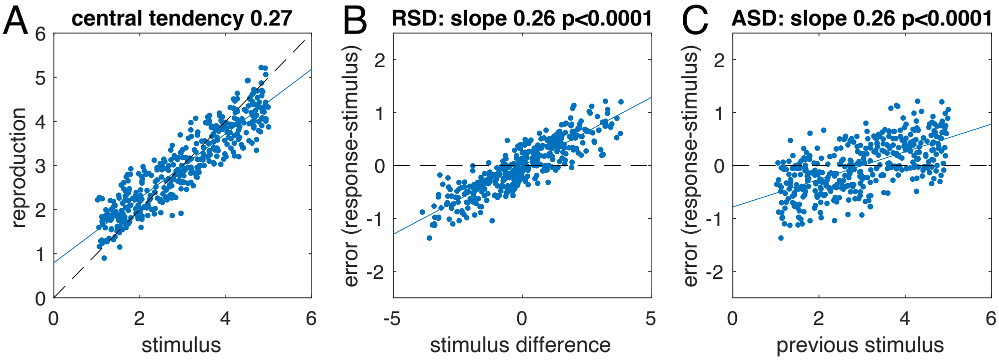


**Figure S10**: Simulation of a simplified version of the serial dependence model by Cicchini et al. (2018) for the same stimulus sequence as in Fig. S2. The slopes of the RSD and ASD are both equal to the central tendency.

The full model of Cicchini et al. (2018) additionally assumes that the weight itself depends on the distance between stimuli (their Eqn. 3.7) so that

$$w_i = 1 - \frac{1}{2 + c(x_{i-1} - x_i)^2}$$

with $c$ being a constant related to the stimulus reliability. This modification of the model, however, does not change the basic relation between central tendency and ASD slope, as exemplified in Fig. S11. In other words, in this model, the serial dependence is equal to the central tendency. Note, however, that according to the weight dependence in the model, the weight $w_i$ can only assume values between 0.5 and 1. Thus, the central tendency according to this model cannot become larger than 0.5. The model thus cannot capture cases such as the "dead salmon" or the noise response shown in Fig. S1, where the response becomes largely independent of the stimulus.

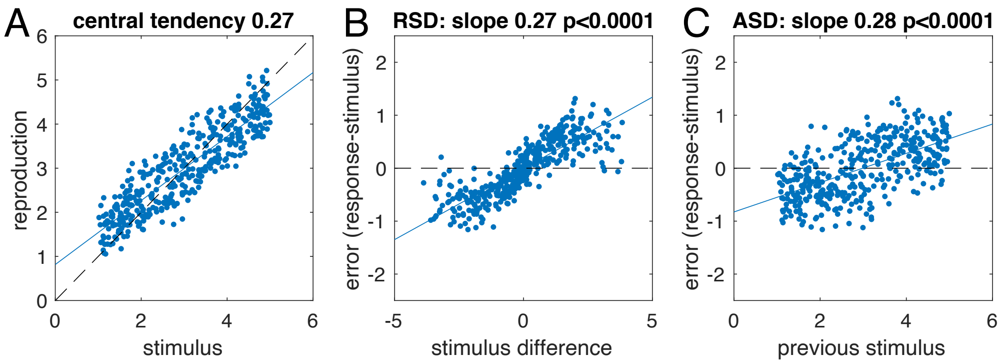


**Figure S11**: Simulation of the full serial dependence model (Cicchini et al. 2018) for the same stimulus sequence as in Figs. S2 and S10. RSD and ASD slopes are again both equal to the central tendency. Due to the dependence of the weight on the stimulus difference the RSD (shown in B) would be better fitted with a polynomial or a difference of Gaussians (as used in Fischer & Whitney 2014). However, a linear fit is still appropriate for the absolute serial dependence (shown in C).

### *References*


Archie, J. P. Mathematical coupling: a common source of error. *Ann. Surg.* **193**, 296–303 (1981)

Bennett, C. M., Miller, M. B. & Wolford, G. L. Neural correlates of interspecies perspective taking in the post-mortem Atlantic Salmon: an argument for multiple comparisons correction. *NeuroImage* **47**, S125 (2009)

Bliss, D. P., Sun, J. J., D'Esposito, M. Serial dependence is absent at the time of perception but increases in visual working memory. *Sci Rep* **7**, 14739 (2017)

Cicchini, G. M., Mikellidou, K., & Burr, D. C. The functional role of serial dependence. *Proc. Biol. Sci.* **285**, 20181722 (2018)

Clifford, C. W. G., Watson, T. L. & White, D. Two sources of bias explain errors in facial age estimation. *R. Soc. Open Sci.* **5**, 180841 (2018)

Curran-Everett, D. Explorations in statistics: correlation. *Adv. Physiol. Educ.* **34**, 186 –191 (2010)

Fischer, J., Whitney, D. Serial dependence in visual perception. *Nat. Neurosci.* **17**, 738-743 (2014)

Glasauer, S. Sequential Bayesian updating as a model for human perception. *Prog. Brain. Res.* **249**, 3-18 (2019)

Glasauer, S. & Shi, Z. Duration reproduction data. G-Node https://doi.org/10.12751/g-node.hdsam3 (2021b)

Griffiths, T. L., Kemp, C. & Tenenbaum, J. B. Bayesian models of cognition. *In: The Cambridge handbook of computational cognitive modeling* (ed. R. Sun) Cambridge University Press (2008)

Holland, M. K. & Lockhead, G. R. Sequential effects in absolute judgments of loudness. *Percept. Psychophys.* **3**, 409–414 (1968).

Kiyonaga, A., Scimeca, J. M., Bliss, D. P. & Whitney D. Serial dependence across perception, attention, and memory. *Trends Cogn. Sci.* **21**, 493-497 (2017)

Petzschner, F. H. & Glasauer, S. Iterative Bayesian estimation as an explanation for range and regression effects: a study on human path integration. *J. Neurosci.* **31**, 17220–17229 (2011)

Petzschner, F. & Glasauer, S. Magnitude estimation data. G-Node https://doi.org/10.12751/g-node.21796b (2020)



### *Acknowledgements*

We thank Mauro Manassi, David Whitney, and Jason Fischer for pointing out that for stimuli on a circular scale the RSD is valid without causing artifacts, and that for the RSD permutation tests should be run as statistical sanity check.
